# Digital Profiling of Tumor Extracellular Vesicle-associated RNAs Directly from Unprocessed Blood Plasma

**DOI:** 10.1101/2024.05.06.592543

**Authors:** Elizabeth Maria Clarissa, Sumit Kumar, Juhee Park, Mamata Karmacharya, In-Jae Oh, Mi-Hyun Kim, Jeong-Seon Ryu, Yoon-Kyoung Cho

## Abstract

Tumor-derived extracellular vesicle (tEV)-associated RNAs hold promise as diagnostic biomarkers, but their clinical use is hindered by the rarity of tEVs among non-tumor EVs. Here, we present EV-CLIP, a highly sensitive droplet-based digital method for profiling EV RNA. EV-CLIP utilizes the fusion of EVs with charged liposomes (CLIPs) in a microfluidic chip. Optimized CLIP surface charge enables exceptional sensitivity and selectivity for EV-derived miRNAs and mRNAs. This approach streamlines detection with minimal plasma volume (20 µL) and eliminates the need for prior EV isolation or RNA preparation, preventing loss of EVs or RNA. In testing with 83 patient samples, EV-CLIP detected EGFR L858R and T790M mutations with high AUC values of 1.0000 and 0.9784, respectively. Its success in serial monitoring during chemotherapy highlights its potential for precise quantification of rare EV subpopulations, facilitating the exploration of single EV RNA content and enhancing understanding of diverse EV populations in various disease states.

**SUMMARY:** EV-CLIP: A highly sensitive digital method for tumor-derived EV RNA detection using undiluted blood plasma samples, promising for precise tumor management.

---

Single extracellular vesicle (EV) analysis is pivotal in biomedical research (*1-3*), offering insights into diseases by examining EV heterogeneity (*4*), identifying disease-specific biomarkers (*5*), and monitoring disease progression through unique nucleic acids (*6-8*), proteins (*9, 10*), or other molecules (*11*). Therefore, investigating single EV contents provides crucial disease process information (*12, 13*). While gold standard quantitative reverse transcription-polymerase chain reaction (RT-qPCR) (*14*) and novel approaches offer exceptional sensitivity (*15*), they still entail laborious steps like EV isolation and RNA extraction (*16, 17*). Additionally, distinguishing RNA signals from tumor-derived EVs (tEVs) and non-tumor EVs remains challenging due to bulk analysis methods (*18, 19*). Consequently, further research is necessary to develop efficient strategies for detecting tEV-originated RNAs.

Membrane fusion, mediated by SNARE proteins is vital for cellular processes, including exocytosis and endocytosis, membrane dynamics, cell division, and signaling(*20, 21*). Extensive research into phospholipid fusion mechanisms (*22*) has led to functional systems with diagnostic and therapeutic potential (*23*). Various EV fusion methods, like pH-dependent (*24*) or polyethylene glycol-mediated (*25*), catechol–metal supramolecular complex (*26*), freeze–thaw-cycle mediated(*27, 28*), and DNA zipper-mediated(*29, 30*), involve diverse molecular compositions facilitating membrane fusion. While previous approaches using fusogenic vesicles (*31*) or aptamer-mediated fusion (*32, 33*) show promise for EV RNA detection, they are complex and time-consuming, limiting clinical utility. Therefore, a generalized, simplified approach is needed for detecting tumor-derived EV (tEV) RNAs in clinical settings.

In this study, we developed a protein-free, charge-induced fusion method to incorporate molecular beacon (MB) loaded charged-liposomes (CLIPs) into EVs. This was achieved in a microfluidic droplet reactor, allowing digital investigation of miRNAs and mRNAs within individual EVs (**Fig. 1a**). By adjusting positive and negative charged lipid ratios, we fine-tuned CLIP surface charge for efficient fusion with EVs. Leveraging microfluidic compartmentalization, our approach enables high-throughput digital profiling of EVs containing target miRNA or mRNA using EV-CLIP fusion reaction (EV-CLIP). In a proof-of-concept study, we detected epidermal growth factor receptor (EGFR) L858R and T790M mutations in blood plasma from 73 patients with lung cancer and 10 healthy donors. Furthermore, EV-CLIP showcased its potential by simplifying the serial monitoring of mutations during chemotherapy, without sample preprocessing, effectively minimizing the risk of EV loss.

**Figure 1.**
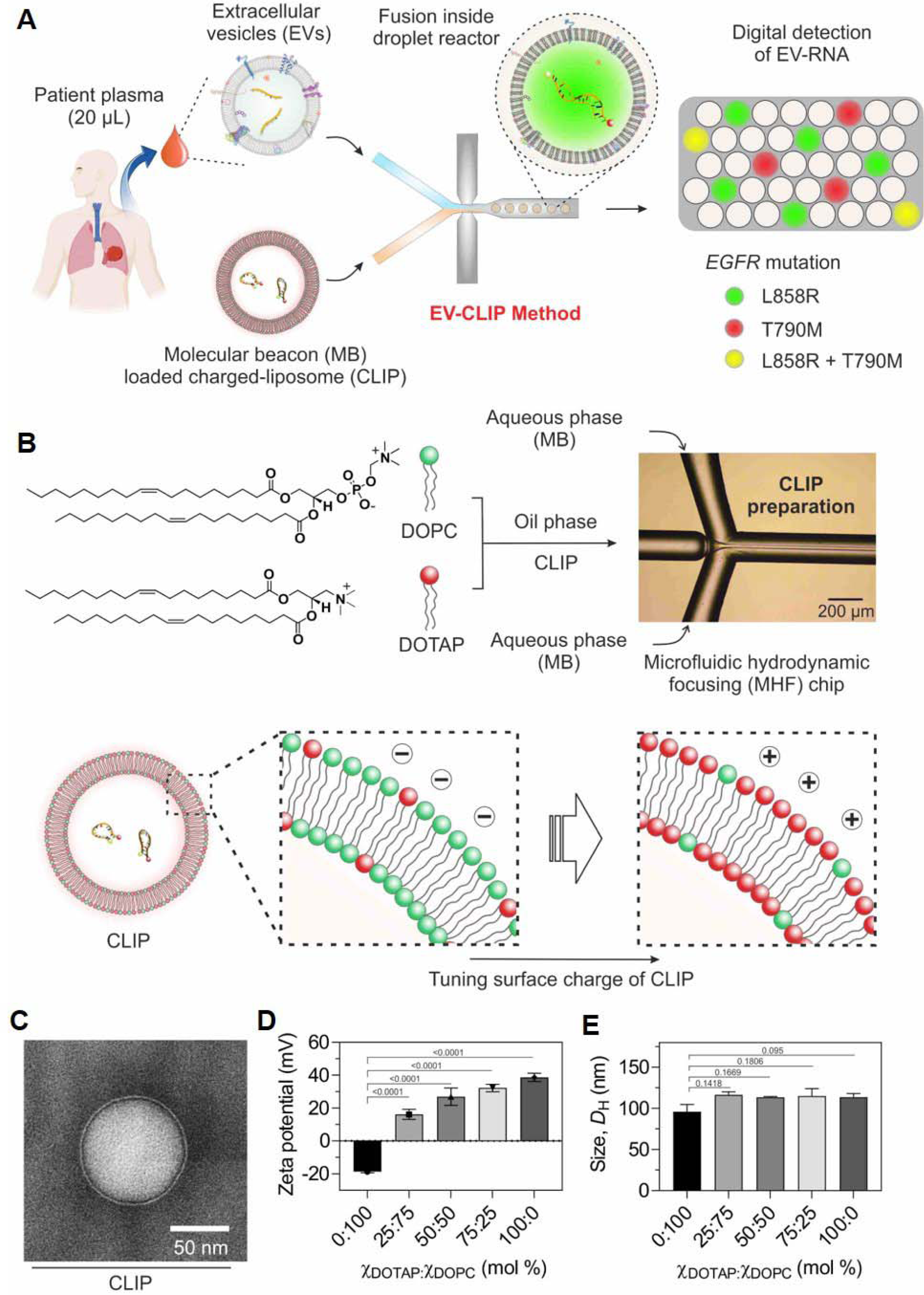
Overview of EV-CLIP method for profiling EV RNA. (**A**) Schematic of the proposed assay, indicating the optical detection of mutations in EV RNA through charge-induced fusion with molecular beacon-loaded charged-liposomes (CLIPs) inside the droplet reactor. (**B**) Schematic of CLIP preparation using microfluidic hydrodynamic focusing method. (**C**) Representative image of the CLIP obtained using a transmission electron microscope (TEM) (diameter: 92.5 ± 5.65 nm, n = 3 with nine data points). (**D**) Zeta potential distribution of CLIPs across varying compositions (mol%) of DOTAP and DOPC (□_DOTAP_: □_DOPC_), presented as mean□±□SD from three independently synthesized CLIP batches. Statistical analysis was conducted using one-way ANOVA with Dunnett’s multiple comparisons test. (**E**) Hydrodynamic size (*D*_H_) distribution of CLIPs across varying compositions (mol%) of DOTAP and DOPC (□_DOTAP_: □_DOPC_), presented as mean□±□SD from three independently synthesized CLIP batches. Statistical analysis was conducted using one-way ANOVA with Dunnett’s T3 multiple comparisons test.

## RESULTS

### Charge-induced EV fusion with CLIPs

Precise control over CLIP charge tuning was achieved by employing two types of lipids: cationic 1,2-dioleoyl-3-trimethylammonium propane (DOTAP) and zwitterionic 1,2-dioleoyl-sn-glycero-3-phosphocholine (DOPC) lipids for CLIP preparation. A microfluidic hydrodynamic focusing (MHF) method (*34*) facilitated efficient and rapid adjustment of CLIP surface charges (**Fig. 1B; Fig. S1 and Video S1**). The microfluidic device directed lipid solutions (DOTAP and DOPC in ethyl alcohol) through the center inlet channel at 5 µL/min, resulting in CLIP formation with an aqueous solution flowing through two lateral inlet channels at 50 µL/min.

CLIP morphology, validated by transmission electron microscopy (TEM), exhibited a diameter of 92.5 ± 5.65 nm, consistent with size measurements obtained via dynamic light scattering (DLS) and nanoparticle tracking analysis (NTA) (**Fig. 1C** and **Fig. S2**). CLIP surface charge was tuned from –18.63 ± 0.76 mV to +38.67 ± 2.51 mV by adjusting lipid composition (DOTAP and DOPC) (**Fig. 1D**). CLIP sizes containing 0, 25, 50, and 75 mol% DOTAP remained stable after 48 h of storage, while those with 100 mol% DOTAP exhibited a consistent 3-fold size increase, reaching approximately 311.50 ± 8.51 nm (**Fig. 1E** and **Fig. S3**).

After optimizing CLIP surface charge, we investigated their fusion with EVs. EVs were isolated from H1975 cells using an efficient, label-free method integrating a lab-on-a-disc system with nanofilters (*35*), enabling rapid enrichment of EVs sized 20–200 nm from cell-culture supernatant (CCS) in just 30 min using a compact centrifugal microfluidic system (**Fig. S4A**).

To ensure digital detection and prevent aggregation, fusion occurred in a droplet reactor (**Fig. S5**). EVs and CLIPs flowed separately through two aqueous inlets at 1.5 µL/min, with an oil phase inlet carrying FC-40 surfactant at 35 µL/min, generating aqueous droplets containing both vesicles when they met at the junction within the oil phase (**Fig. 2A** and **Video S2**). Controlling the number of EVs and CLIPs in each droplet was feasible by adjusting the initial input concentration using Poisson distribution calculation (**Fig. S6**). We controlled the input concentration of EVs (10–10^5^ EVs/µL) and CLIPs (10^4^–10^6^ CLIPs/µL) within a 27.40 ± 2.06 µm-sized droplet reactor (**Fig. S7**).

**Figure 2.**
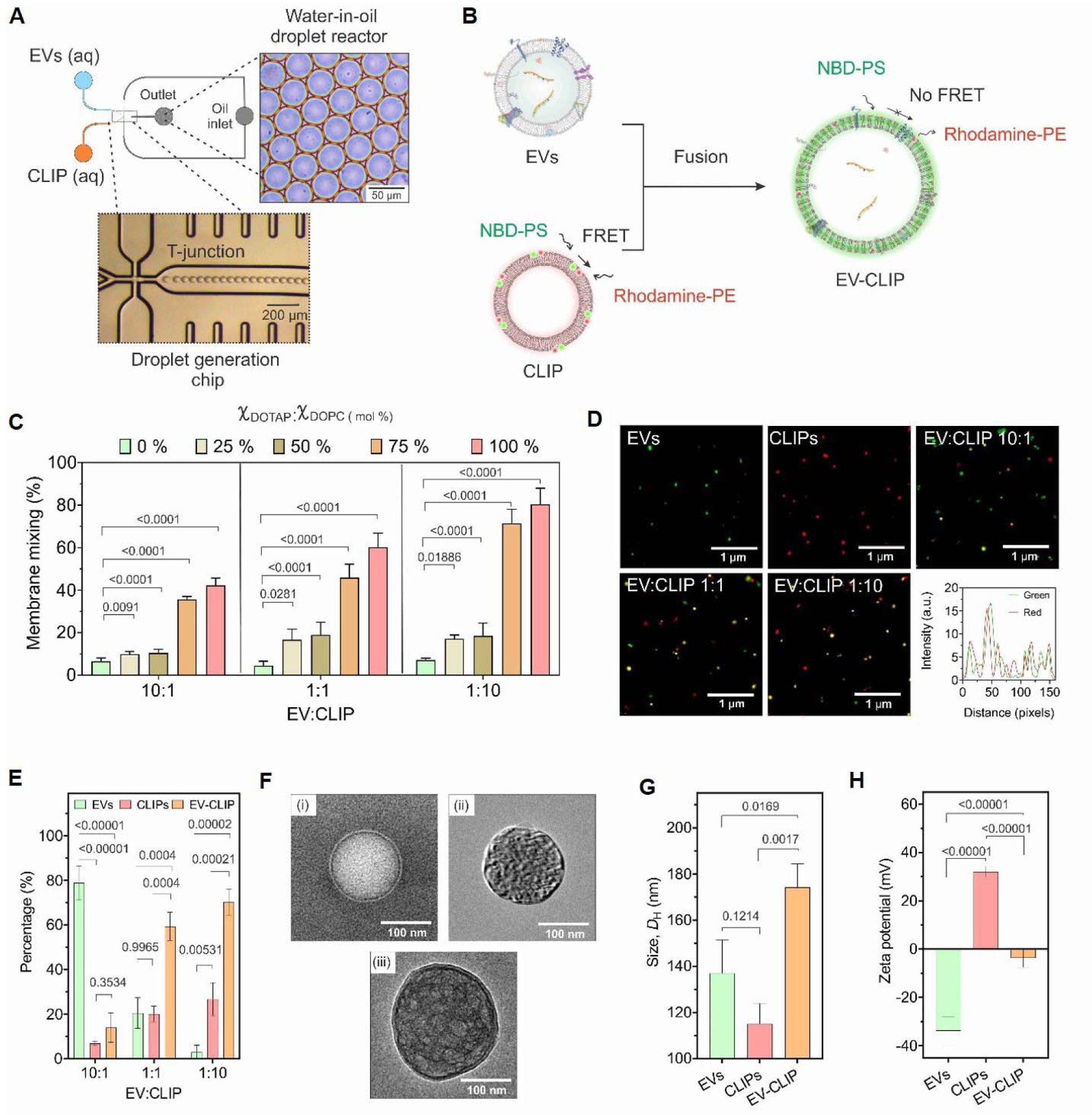
Controlled fusion of EVs and CLIPs. (**A**) Schematic of CLIP and EV fusion using the droplet generation chip. The microfluidic device was designed for generating water-in-oil droplet reactors at the flow-focusing junction, where two aqueous phases (EVs and CLIPs) met and introduced into the oil stream (surfactant PFPE-PEG in FC-40). (**B**) Schematic illustration of the Förster resonance energy transfer (FRET)-based lipid mixing assay to investigate exosome fusion event. The average distance between the FRET-pair donor (green, nitrobenzoxadiazole) and acceptor (red, rhodamine) lipid probe increases following charged-based fusion of a labeled CLIP membrane and unlabeled EV membrane, resulting in decreased efficiency of FRET (represented by straight black arrow). PE, phosphoethanolamine; PS, phosphatidylserine. (**C**) Membrane mixing percentage of CLIPs: EVs in different ratios with various CLIPs surface charges after 30 min. Data represent mean ± SD, n = 3 independent membrane mixing experiments. All statistical analyses were conducted using one-way ANOVA using Dunnett’s multiple comparisons test. (**D**) High-resolution confocal laser scanning microscope (CLSM) images of EVs, CLIPs, and fused vesicles (EV-CLIP) in three different ratios of EVs and CLIPs: 10:1, 1:1, and 1:10. Intensity analysis shows the colocalization of EVs (green) and CLIPs (red). (**E**) Population of each vesicle type at different ratios was analyzed from 10 independent images, with vesicle counts ranging from 100 to 160. Statistical analysis was performed using one-way ANOVA using Tukey’s multiple comparison test. (**F**) Representative image of CLIPs, EVs, and fused vesicles obtained using the TEM (core diameter of EV-CLIP: 174.85 ± 13.83 nm, n = 5). Scale bar: 100 nm. (**G**) Vesicle size before and after controlled fusion (EVs: 137.03 ± 14.30 nm, CLIPs: 115.03 ± 8.85 nm, EV-CLIPs: 174.10 ± 10.28 nm). Data represent mean ± SD, n = 3. All statistical analyses were performed using one-way ANOVA using Tukey’s multiple comparison test. (**H**) Zeta potential of the vesicles before and after controlled fusion. Data represent mean ± SD, n = 3. Statistical analysis was conducted using one-way ANOVA using Tukey’s multiple comparison test.

Membrane mixing efficiency was evaluated using a Förster resonance energy transfer (FRET)- based lipid mixing assay to optimize fusion conditions (*36*). CLIPs were dual-labeled with nitro-2,1,3-benzoxadiazole-4-yl (NBD) as the donor and Lissamine rhodamine B (Rho) as the acceptor (*37*), facilitating energy transfer from the excited NBD donor to the Rho acceptor when they were in proximity (**Fig. 2B**). Dual labeling showed no significant effect on surface charge (**Fig. S8**). Testing EV (10^5^ EV/µL) fusion with CLIPs in different molar ratios (10:1, 1:1, and 1:10) revealed increased membrane mixing efficiency corresponding to DOTAP percentage (6.6 ± 1.5%–42.3 ± 3.5%, 4.6 ± 2.0%–60.2 ± 6.5%, and 7.2 ± 0.9%–80.2 ± 7.6%, respectively) (**Fig. 2C**). However, CLIPs with 100 mol% DOTAP exhibited instability and aggregation over time, despite achieving the highest fusion efficiency (80.2 ± 7.6%) (**Fig. S3**). Comparing EV-CLIP fusion in the droplet reactor with bulk-scale experiments revealed aggregations in the latter (**Fig. S9**).

To visually demonstrate EV-CLIP fusion, we labeled EVs with green fluorescence dye and CLIPs with red fluorescence dye, then imaged them using confocal laser scanning microscopy (CLSM). Testing different EV to CLIP ratios (10:1, 1:1, and 1:10), we observed varying populations of each vesicle type. With the 10:1 ratio, EVs outnumbered CLIPs and EV-CLIPs; with the 1:1 ratio, EVs and CLIPs were nearly equal, while in the 1:10 ratio, more CLIPs and fused vesicles were observed. CLSM imaging confirmed dye colocalization in fused vesicles (**Fig. 2D**). Quantifying Pearson’s correlation coefficient (PCC) showed highest fusion efficiency at a 1:10 EV to CLIP ratio, yielding a PCC of 0.69 ± 0.09 (**Fig. 2E** and **Fig. S10**).

TEM analysis confirmed spherical morphology of individual and fused vesicles (**Fig. 2F**) without membrane breakage. DLS showed increased vesicle size after fusion (**Fig. 2G**). Increasing DOTAP percentage in CLIPs (10^6^ CLIP/µL) for fusion with EVs (10^5^ EV/µL) maintained stable vesicle size until 75 mol% DOTAP; 100 mol% DOTAP substantially increased size (**Fig. S11A**). Zeta sizer analysis validated fusion by observing increased zeta potential with higher DOTAP percentage (**Fig. 2H** and **Fig. S11B**). Long-term stability assessment over 48 h confirmed stability of fused vesicles, CLIPs, and EVs (**Fig. S12**). These findings demonstrate efficient EV-CLIP fusion. Hence, we chose 75 mol% DOTAP and 10^6^ CLIP/µL for subsequent experiments to maximize fusion, ensuring each droplet contains one or more CLIPs (**Fig. S6**).

### Digital detection of tEV-associated miRNAs using EV-CLIP

After demonstrating efficient fusion between CLIPs and EVs, we evaluated the fluorescence response of CLIP-encapsulated MB to miRNA inside tEVs using a microfluidic cascade platform. This tested the effectiveness of the charge-induced fusion strategy for detecting EV RNA in their natural environment. An MB with a cyanine3 fluorophore at the 5’ end, a tetrachlorofluorescein quencher at the 3’ end, and complementary bases at both ends (*38*), facilitating proximity-induced fluorescence quenching, served as the detection probe (**Table S1**). tEVs were initially detected using an MB targeting microRNA-21 or miR-21, known to be upregulated in various tumors (*39*). During CLIP generation, miR-21-specific MB (miR-21 MB) was introduced via the aqueous phase, with concentrations ranging from 1 fM to 2 µM, optimizing fluorescence intensity. The optimal signal was achieved at 1 µM of miR-21 MB (**Fig. S13**), with no notable impact on CLIP size or zeta potential (**Fig. S14**).

To demonstrate the feasibility of our approach, we detected H1975 lung cancer cell line-derived EVs in phosphate-buffered saline (PBS) by fusing them with CLIPs containing miR-21 MB using the microfluidic cascade platform. This system integrates CLIPs produced by the MHF-chip with a droplet generation chip and a detection chamber chips for imaging, minimizing sample loss (**Fig. 3A–B**). Maintaining a constant CLIP concentration (10^6^ CLIPs/µL) ensured each droplet contained one or more CLIPs. Varying H1975 EV concentration (10–10^5^ EVs/µL) proportionally increased the number of droplets with fluorescent signal (**Fig. 3C)**. Imaging conditions and signal processing were optimized to visualize EVs within the droplets (**Fig. S15**, **Fig. S16**). The EV-CLIP system could detect miR-21 positive EVs within a range of 10 to 10^5^ EVs/µL, with a limit-of-detection (LOD) of 79 EVs/µL (**Fig. 3D–E**). Control experiments without EVs, only free miR-21, showed no signal, confirming system specificity (**Fig. S17**). Moreover, the system distinguished miR-21 levels in EVs from healthy donors and cancer patients’ plasma (**Fig. 3F** and **Fig. S18**), indicating higher miR-21 expression in cancer patients, consistent with literature (*40*). Here, plasma was used directly without prior EV isolation step.

**Figure 3.**
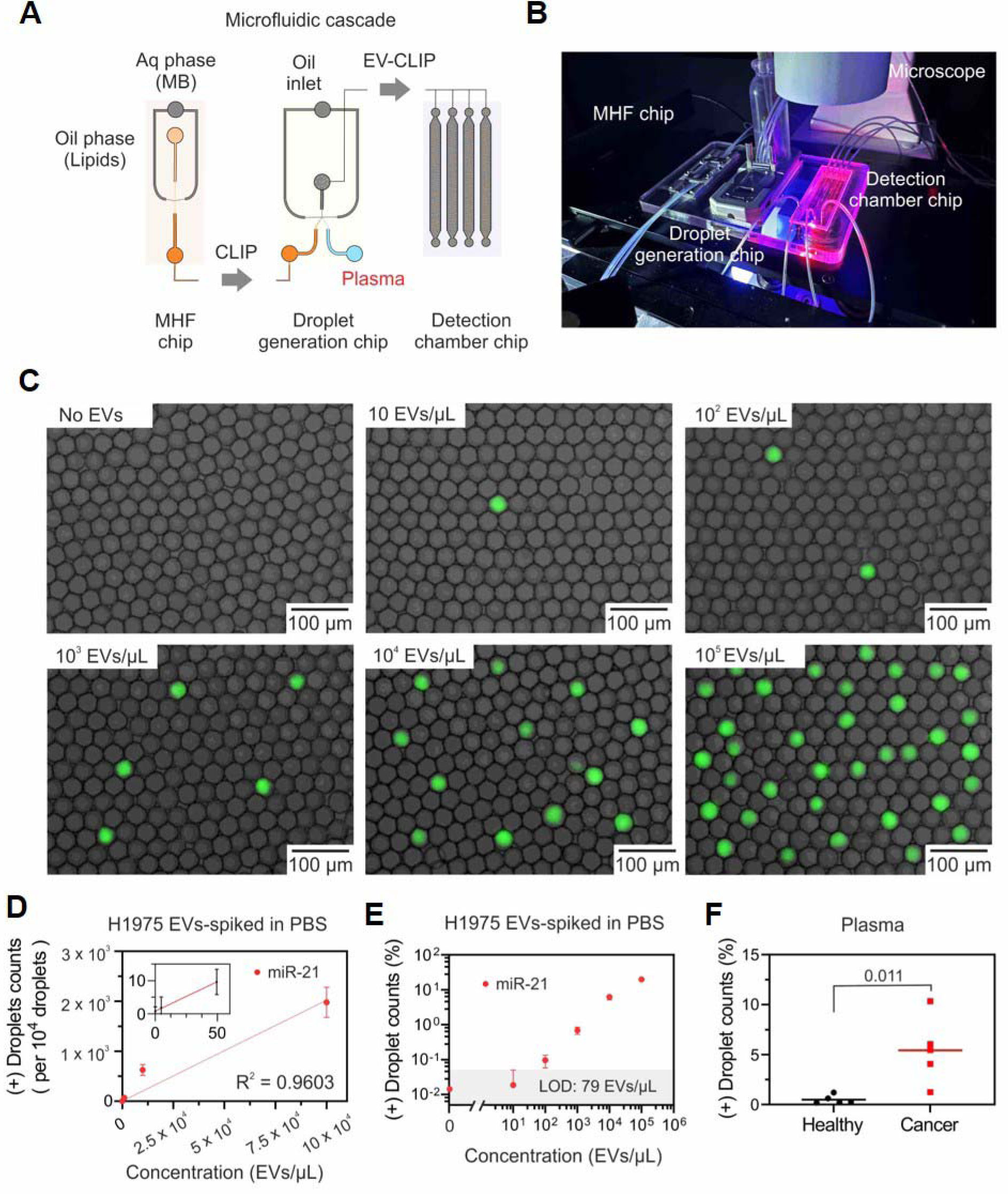
Digital detection of tumor-derived EV miR-21 using the microfluidic cascade platform. (**A**) Schematic of the microfluidic cascade-based detection starting from CLIP preparation using the MHF chip, clinical plasma samples into the droplet generation chip, and RNA profiling in the detection chamber chip using CSLM imaging. (**B**) Microfluidic cascade setup. (**C**) Representative fluorescent images of miR-21 molecular beacon-based detection of various concentrations of H1975 EVs in phosphate-buffered saline (PBS). (**D**) miR-21 detection with respect to H1975 EV concentration on a linear scale. Data represents the mean ± SD from n = 3 experiments, with a limit of detection (LOD) of 79 EVs/µL. (**E**) miR-21 detection with respect to H1975 EV concentration on a logarithmic scale. (**F**) Comparison of miR-21-positive droplets in 5 healthy donors and 5 patients with cancer. Data represents the mean ± SD from n = 3 experiments. Statistical analysis was performed using a two-tailed unpaired Student’s *t*-test.

### Digital profiling of EGFR mutations using tEV-derived mRNAs

We further evaluated the EV-CLIP method by detecting EGFR L858R and T790M mutations in H1975 EVs, known to carry these mutations (*41*). EGFR mutation-specific MB sequences (*42*) were introduced into CLIPs, similar to miR-21 MB loading. The inclusion of L858R and T790M MBs did not alter CLIP size or zeta potential significantly (**Fig. S14**). Under the same experimental conditions as miRNA detection, the only difference being with specific MBs targeting EGFR mutations, we successfully detected these mutations. The fluorescent signal was observed exclusively in the presence of target EVs, with an LOD of 1348 EVs/µL for L858R and 3595 EVs/µL for T790M (**Fig. 4A–B**). This LOD was approximately 17 and 46 times higher than that for miR-21, suggesting lower mRNA abundance in EVs. Before analyzing cancer patient plasma samples, we validated mutation detection by spiking varying concentrations of H1975-derived EVs into healthy human plasma (**Fig. 4C–D**). The concentrations ranged from 10 to 10^5^ EVs/µL, and the detection results for both mutations were aligned with those obtained for EVs spiked in PBS (**Fig. 4E**), confirming the robustness of our EV-CLIP method.

**Figure 4.**
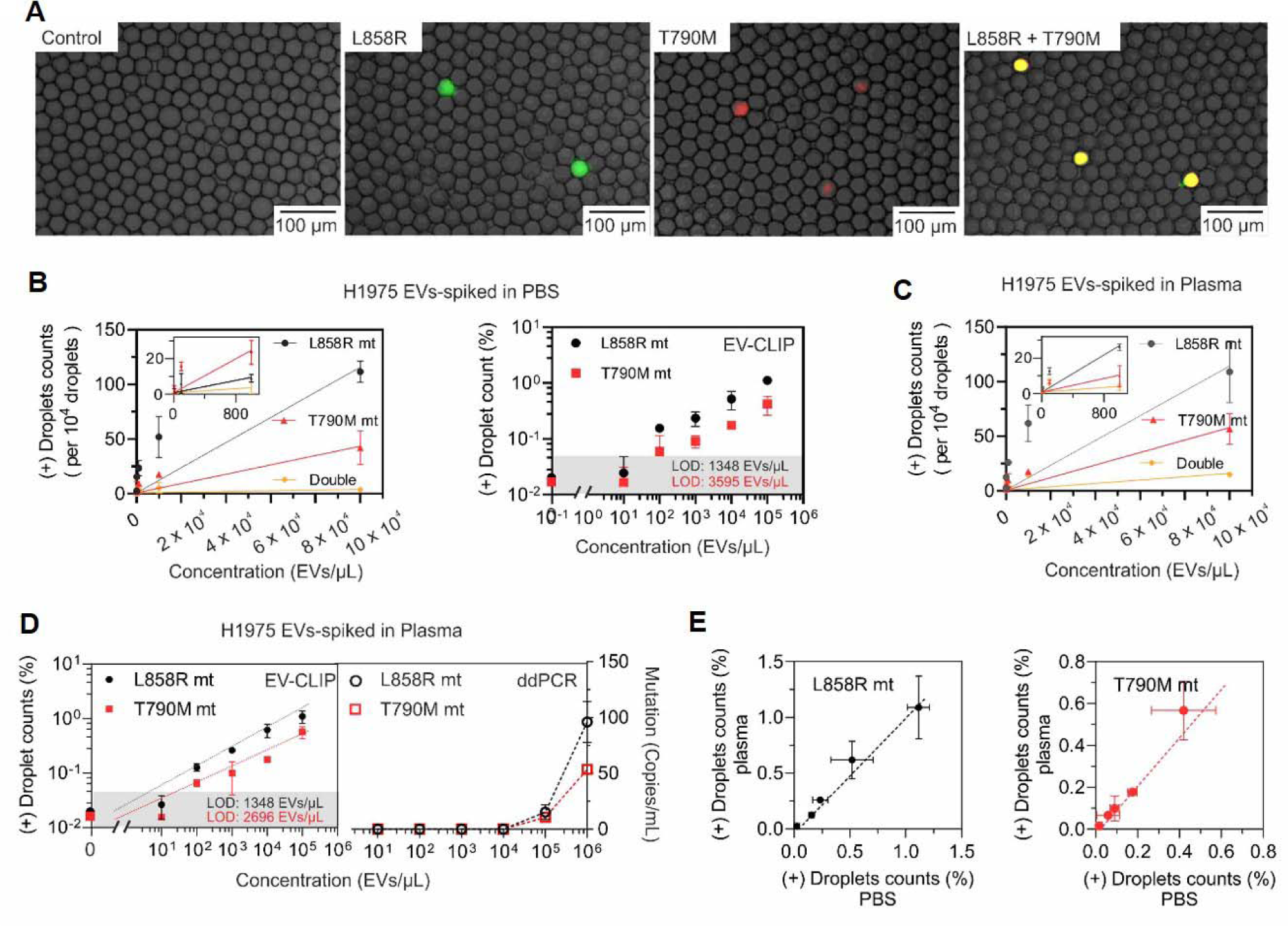
Digital detection of tumor-derived EV EGFR L858R and T790M mRNAs. (**A**) Representative CLSM images of mutation detection (from left to right: negative control, EGFR L858R mutation, EGFR T790M mutation, and both EGFR L858R and T790M mutations) (λ_ex_□=□488 and 375Lnm). Scale bar = 100 μm. EGFR L858R, T790M, and double (both EGFR L858R and T790M) mutation detection with respect to the concentration of H1975 EVs in (**B**) PBS, (**C**) human plasma on a linear scale, and (**D**) human plasma on a logarithmic scale. A gold standard, droplet digital polymerase chain reaction (ddPCR) was conducted to detect the presence of mutations in EV samples spiked in human plasma (right panel). (**E**) Correlation of detection readout of EGFR L858R and T790M mutations in PBS and human plasma. Data represents the mean ± SD from n = 3 independent analyses.

EVs, containing miRNA and mRNA, have emerged as potential intercellular communication vehicles and less invasive disease biomarkers (*43*). However, accurately quantifying these molecules within EVs is challenging. Many studies have used excessive EVs and necessitated complex sample preparation steps like EV separation and RNA extraction, potentially introducing variability in quantitative estimates. The relative abundance of specific miRNA or mRNA may vary depending on RNA or cell types, with reports indicating that fewer than one copy of miRNA (10^-5^–10^-1^) is typically packaged per EV (*44*). Herein, we did not count individual EVs but quantified RNA by measuring the number of droplets containing specific EVs. We gained valuable insights by estimating stoichiometry through the fraction of specific marker-positive droplets normalized by the fraction of droplets carrying a single EV. In this digital detection approach with 1000 EVs/µL, 99% of the droplets were empty, and 1% were with 1 EV according to the Poisson distribution calculation. We observed that 0.69% of the droplets carried miR-21, suggesting that roughly 69% of H1975 EVs carry miR-21. Additionally, approximately 24% of EVs from H1975 culture may contain L858R-positive EGFR mRNA, 9% T790M-positive, and 4% both mutations.

Our EV-CLIP method could detect single EV per nanoliter, making it approximately 100 times more sensitive than standard ddPCR-based mRNA quantification (LOD = 1,348 EVs/µL for L858R using EV-CLIP method and 100,000 EVs/µL using ddPCR) (**Fig. 4D**). Compared to previous methods for miRNA (**Table S2**) and mRNA (**Table S3**) detection, our EV-CLIP method offers equivalent or superior sensitivity. It requires less than 20 µL of plasma, eliminating the need for time-consuming and complicated sample preparation processes such as EV separation, RNA extraction, and amplification. This enables rapid and sensitive tEV miRNA or mRNA detection. To the best of our knowledge, the EV-CLIP method represents the first platform for the digital detection of EV-derived RNAs, distinguishing it from previous digital methods primarily focused on detecting EV proteins (**Table S4**).

### EV-CLIP detected EGFR mutation in lung cancer plasma

To assess its clinical utility as companion diagnostics, we focused on EGFR L858R and T790M mutations in clinical samples obtained from patients with lung cancer (**Fig. 5A**). Lung cancers with EGFR gene mutations initially respond to tyrosine kinase inhibitors (TKIs). However, drug resistance, often caused by the T790M mutation, limits TKI effectiveness. Although liquid biopsies for EGFR mutation detection exist, they exhibit lower sensitivity than tissue rebiopsies (*45, 46*). In this context, we investigated EV-based nucleic acid detection as a promising clinical diagnostic approach.

**Figure 5.**
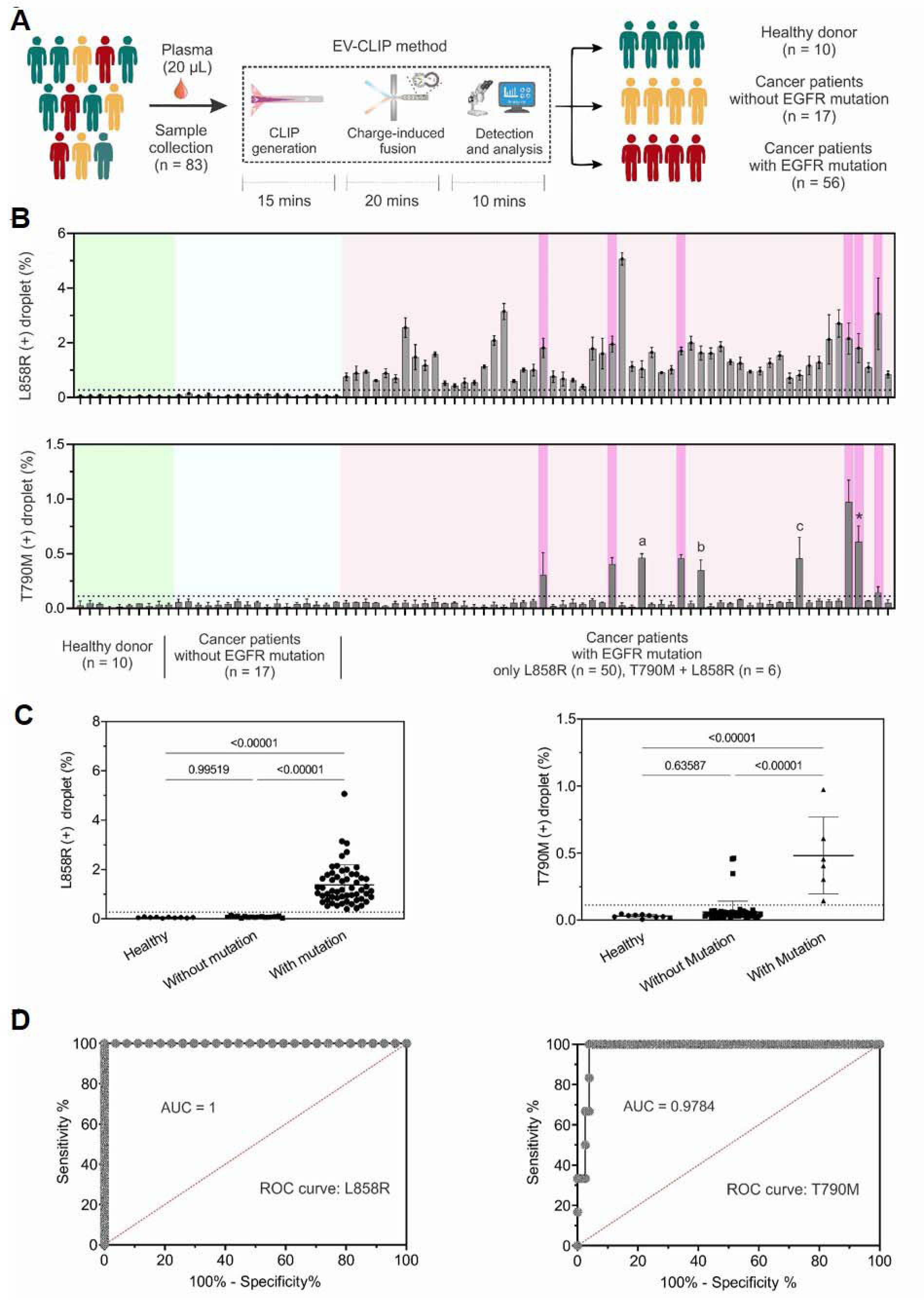
Detection of EGFR L858R and T790M mutation from lung cancer samples using the EV-CLIP method. (**A**) Schematic illustration of the mutation detection using EV-CLIP method. (**B**) Signal quantification of EGFR L858R and T790M detection from 10 healthy donor samples and 73 lung cancer samples. The green area represents healthy donors; blue indicates patients without the mutation; pink indicates patients with EGFR L858R mutation, with the dark pink area representing patients with both L858R and T790M mutations. For patient “*,” the T790M mutation was detected when it was still undetectable in tissue samples using the conventional EGFR Mutyper method, later confirmed with bronchoalveolar lavage fluid (BALF). Data represents mean ± SD from n = 3 independent analyses. (**C**) Comparison of positive (+) droplet counts between 10 healthy donors and 73 patients with lung cancer for L858R and T790M mutation detection. Data represents the mean ± SD from n = 3 independent analyses. Statistical analyses were conducted using one-way ANOVA using Tukey’s multiple comparison test. (**D**) Receiver operating characteristic (ROC) curve for L858R and T790M mutation detection, with the area under the curve (AUC) being 1.00 and 0.9784, respectively. Each data point represents the average from three independent repetitions.

Our study included blood plasma samples from a cohort of 10 healthy donors and 73 patients with lung cancer drawn from various hospitals (**Table S5**). Among them, 17 lung cancer samples exhibited no EGFR mutations, whereas 56 showed the presence of EGFR L858R mutations, with 6 samples harboring both EGFR L858R and T790M mutations. The distribution of samples per mutation type may not statistically represent the broader population of such mutations, as we intentionally included a higher number of samples with positive EGFR mutation. Samples were directly analyzed without preprocessing, revealing higher mutation-positive droplet counts in cancer patients’ plasma (**Fig. 5B** and **Fig. S25**). Patients with mutations showed significantly more mutation-positive droplets than those without mutations (**Fig. 5C** and **Fig. S19-S24**). This direct detection from plasma, without complex preprocessing, demonstrates the potential of EV-CLIP as a sensitive and efficient tool for EGFR mutation screening in lung cancer patients.

Remarkably, the EGFR L858R mutations could be detected with 100% accuracy in both early-stage (n=24) and advanced-stage (n=32) patients, irrespective of cancer stage. Additionally, our EV-CLIP platform revealed a potential correlation between the percentage of L858R-positive droplets and cancer stage (**Fig. S26**), consistent with literature suggesting a link between mRNA levels and cancer staging (*47*). Notably, our EV-CLIP method, using just 20 µL of plasma, achieved 100% accuracy in detecting L858R mutations in early-stage lung cancer patients (**Fig. S27**). This represents a substantial improvement, particularly for early-stage patients with low mutant allele frequencies, over FDA-approved blood-based tests relying on qPCR, ddPCR, or NGS, which analyze cell-free DNA (cfDNA) in peripheral blood (*48*). Notably, the mutation rate remained consistent across different stages, aligning with existing literature reporting similar EGFR mutation rates between earlyLstage and advancedLstage groups (*49, 50*).

The platform exhibited highly sensitive and specific detection of EGFR L858R (1.000 AUC, 100% sensitivity, 100% specificity, and 100% accuracy) and T790M (0.978 AUC, 100% sensitivity, 96.10% specificity, and 96.38% accuracy) mutations, as shown by the ROC curves (**Fig. 5D**). Notably, T790M mutations were identified in samples from 6 patients (**Fig. 5B**), all with confirmed tissue-verified mutations. One patient labeled “*” exhibited an early T790M mutation, initially undetectable in tissue or plasma-derived ctDNA analysis. However, erlotinib resistance was subsequently confirmed, and the T790M mutation was verified through bronchoalveolar lavage fluid (BALF)-based DNA analysis. Furthermore, the platform detected T790M mutations in three other patient samples denoted as a, b, and c, where mutations were not tissue-confirmed, categorizing them as “false positives”. Resistance cross-validation remains challenging due to patient-specific conditions. Patient “a” received osimertinib treatment for two months, patient “b” developed dacomitinib resistance after 21 months, confirmed through an EGFR-T790M mutation identified in a bone biopsy, and patient “c” experienced brain metastasis 6 months post-afatinib treatment and, unfortunately, passed away, with limited details available from the hospital.

To further evaluate the clinical utility of our platform, we examined the presence of EGFR L858R and T790M mutations over time in samples from patients with lung cancer undergoing treatment. The cohort comprised 33 samples from 4 patients, collected at various stages during treatment with EGFR-targeting drugs such as gefitinib and osimertinib.

Patient 1, a 57-year-old female with stage IV non-small cell lung cancer, initially responded to gefitinib treatment, which reduced the size of her primary tumor. However, after 8 months, drug resistance emerged alongside worsening pleural metastasis. Subsequent treatment with osimertinib, prompted by plasma EGFR T790M detection using the Cobas® EGFR Mutation Test v2, lasted 9 months. Our EV-CLIP-based detection corresponded with tissue biopsy results, detecting an increase in T790M-positive droplets before confirmation (**Fig. 6A, Fig. S28A, S29A**).

**Figure 6.**
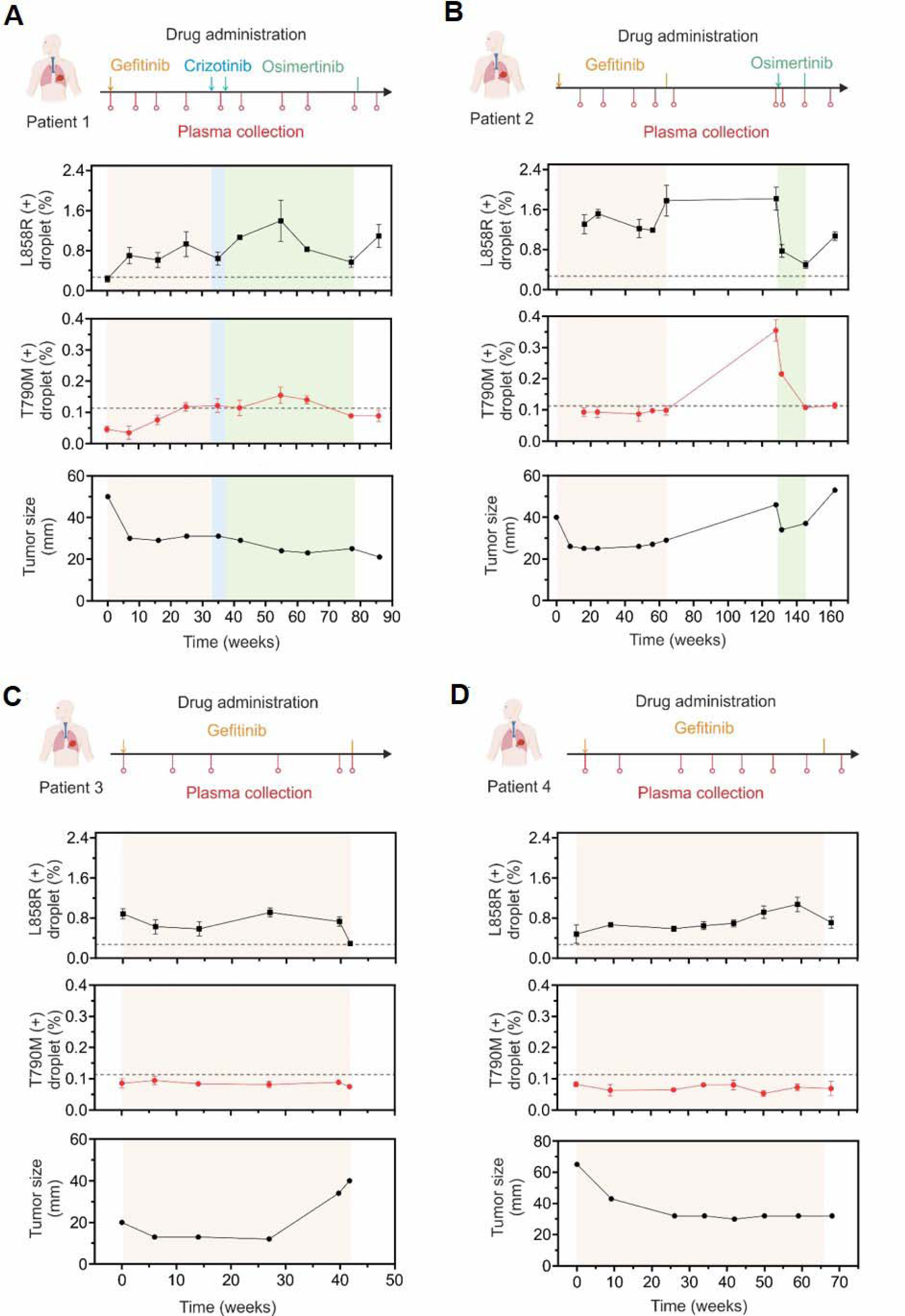
Monitoring of EGFR L858R and T790M mutation from lung cancer patient samples using the EV-CLIP method. (**A**) L858R and T790M mutation detection for patient 1. Area colored in orange represents gefitinib treatment, blue represents crizotinib treatment and green represents osimertinib treatment, corresponding to the sampling and treatment timeline shown on the schematic above the result. The primary tumor size of patient 1 decreased after gefitinib treatment, and T790M-positive droplet percentage increased before osimertinib treatment. (**B**) L858R and T790M mutation detection for patient 2. Area colored in orange represents gefitinib treatment, and green represents osimertinib treatment. The T790M -positive droplet percentage was elevated during the treatment absence after gefitinib resistance occurred. The T790M and L858R droplet percentage decreased after osimertinib treatment. (**C**) L858R and T790M mutation detection for patient 3. Area colored in orange represents gefitinib treatment. The tumor size increased after 10 months of treatment, however neither L858R nor T790M droplet percentage showed significant difference. (**D**) L858R and T790M mutation detection for patient 4. Area colored in orange represents gefitinib treatment. The tumor size decreased after gefitinib treatment, and the L858R droplet percentage did not show significant difference during gefitinib treatment. T790M droplet percentage remained under the detection range throughout whole treatment period. All data points represent the mean ± SD from n = 3 independent analyses.

Patient 2, a 52-year-old female with stage IV adenocarcinoma and bone and pleural metastasis, initially responded to gefitinib for 15 months. Following disease progression, plasma EGFR T790M detection led to osimertinib treatment for 4 months. Again, our EV-CLIP platform detected an elevation in T790M mutation before confirmation (**Fig. 6B, Fig. S28B, S29B**).

Patients 3 and 4, also non-smoking females with EGFR L858R mutation-positive tumors, exhibited partial responses to gefitinib for 10 and 15 months, respectively. However, no EGFR T790M mutation was detected in plasma, pleural fluid, or bronchoalveolar lavage fluid. This suggests alternative resistance mechanisms. EV-CLIP did not detect T790M mutation in these patients (**Fig. 6C, D, Fig. S28C-D, S29C-D**).

## DISCUSSION

We developed a simple, robust, and generalizable method for digital profiling of EV RNA, incorporating an amplification-free, extraction-free approach that eliminates the necessity for conventional steps of EV isolation, lysis, RNA extraction, and RNA analysis using nucleic acid amplification. Leveraging the advantages of high fusion rates, rapid applicability, and broad usability, EV-CLIP demonstrates remarkable sensitivity and selectivity in detecting tEV miRNAs and mRNAs without the need for sample lysis, thereby streamlining the detection process and preserving EV integrity. This method eliminates the requirement for sample preprocessing, simplifying the detection process while preventing EV loss. Overall, the EV-CLIP method shows great promise in the clinical setting, offering precise quantification of rare EV subpopulations and opening new avenues to explore fundamental questions in cancer biology. Beyond its direct applications in diagnostics, this fusion system exhibits substantial potential for diverse implementations, including the development of drug delivery carriers, profiling EV markers, and numerous other possibilities. While the proof-of-concept study highlights the translational potential of EV RNA analysis in clinical samples, the clinical applicability for cancer diagnosis necessitates additional tests with larger cohorts. To enhance its utility and reliability in hospital settings, we anticipate the need for further refinement through the incorporation of automation and multi-marker analysis. This advancement is crucial for improving our comprehension of EV biology and their roles as biomarkers and molecular messengers in cell-to-cell communications.

## MATERIALS AND METHODS

### Study Design

The primary objective of this study was to develop a method for detecting tumor-derived EV RNA through charge-based fusion inside a droplet. The study encompassed the generation of CLIP and microfluidic-aided EV-CLIP fusion within a droplet generator. To evaluate the platform’s effectiveness, it was employed to detect EGFR L858R and T790M mutations in a total of 83 human plasma samples consisting of 10 healthy donor, 17 patients with lung cancer without EGFR mutation, and 56 patients with lung cancer expressing EGFR mutation, in which 6 patients exhibited both L858R and T790M mutation. Furthermore, the platform was subjected to additional testing to monitor these mutations in 33 samples obtained from 4 patients with lung cancer undergoing treatment

### CLIP synthesis

CLIPs were synthesized using the MHF method on a commercialized platform (Dolomite Microfluidics, Royston, UK). Briefly, the lipid mix for the CLIPs was created by dissolving various ratios of 18:1 TAP or DOTAP (1,2-dioleoyl-3-trimethylammonium propane, Avanti 890890P) and 18:1 (Δ9-Cis) PC or DOPC (1,2-dioleoyl-sn-glycero-3-phosphocholine, Avanti 850375P): 0, 25, 50, 75, and 100 mol% DOTAP in 1 mL pure ethanol (UN1170; Duksan Pure Chemicals, Ansan, South Korea) filtered using 0.2 µm syringe filter (Minisart^®^ NML Syringe Filters, S6534) to reach a final concentration of 5 mg/mL. CLIPs were synthesized using a glass microfluidic device (5 Input Chip 3D 150 µm (Part No. 3200834), Dolomite Microfluidics) at an aqueous phase flow rate of 50 µL/min (1 × PBS, pH 7.4; Gibco) and an oil phase (lipid mix) flow rate of 5 µL/min. The suspension was collected for 15 min in a 1.5 mL microcentrifuge tube and analyzed using a NanoSight instrument (Nanosight NS500; Malvern Panalytical, Worcestershire, UK) and Malvern Zetasizer (Nano ZS) to determine the concentration, size, and zeta potential distribution. During the automated detection, the final lipid concentration was adjusted to 3 mg/mL to yield a CLIP concentration of 10^9^ particles/mL.

### Cell Culture

H1975 cells were obtained from American Type Culture Collection (ATCC) and seeded at a density of 1×10^6^ cells per 10-cm culture dishes at 37 °C in the presence of 5% CO_2_ in Dulbecco’s Modified Eagle Medium (DMEM) (11965092; Gibco) supplemented with 5% (v/v) FBS (Gibco) and 1% antibiotic/antimycotic (100 U/mL penicillin and 100 mg/mL streptomycin, 15240062; Gibco) in static condition. After 24 h, the cells were washed twice with 1 × PBS (pH 7.4, Gibco) to remove any debris, and were left to grow until they reached approximately 80% confluency. After being passaged two times, the cells were washed with 1 × PBS (pH 7.4, Gibco) and then cultured in DMEM (Gibco, 11965092) supplemented with 5% EV-depleted FBS (Systems Biosciences Inc) and 1% antibiotic/antimycotic for 48 h for EVs isolation.

### Extracellular vesicles (EVs) isolation from cell culture

EVs were isolated from cell culture media using a commercialized ExoDisc platform, a centrifugal disc equipped with anodized aluminum oxide filters with a pore diameter of 0.02 µm (Exodisc^TM^-C, LabSpinner)(*35*). Briefly, 70–80% confluent 10-cm culture dishes of cells were washed twice with 1 × PBS (pH 7.4, Gibco) and cultured in their respective medium supplemented with 5% EV-depleted FBS (Systems Biosciences) and 1% antibiotic/antimycotic for 48 h. Subsequently, the cell culture supernatant was collected and centrifuged at 300 ×*g* for 10 min and 2000 ×*g* at 4°C for another 10 min to remove cell debris, then passed through a 0.2 µm syringe filter (S6534, Minisart^®^ NML Syringe Filters). The clarified supernatant was concentrated using the ExoDisc platform and passed through the filter by centrifugation at 3000 rpm (approximately 500 ×*g*). Further, 100 µL of concentrated EVs from the collection chamber was resuspended in 1 × PBS with a dilution factor of two. The isolated EVs were then analyzed using a NanoSight instrument to determine the concentration and size and Malvern Zetasizer to determine zeta potential distribution. The EVs were aliquoted and stored at -80 °C for further experiment.

### EV-CLIP fusion

The EV-CLIP fusion was conducted using a commercialized platform (Dolomite Microfluidics). The glass chip (3200529, Droplet generation chip; Dolomite Microfluidics) comprises four channels: two aqueous phase channels, one oil phase channel, and one output channel. Briefly, equal concentrations of CLIPs and EVs in PBS were loaded into the aqueous channels of the microfluidic chip, where 30 µm-diameter droplets were generated in the FC-40 (RAN Biosciences, 008-FluoroSurfactant-2wtF-50G) oil phase. The number of EVs and CLIPs can be controlled by varying the initial sample input concentration, ranging from 10 to 10^5^ particles/µL. The pressures of the pumps was set to 2500 and 1100 mBar for oil and aqueous phases, respectively. The droplets were collected for 20 min in microcentrifuge tubes and stored at 4 °C for further experiments.

### Transmission electron microscope

The vesicle suspension (2.5 µL, 3 × 10^9^ particles mL^-1^) of the respective solutions (CLIPs, EVs, and EV-CLIPs) was vortexed for 30 s and drop-casted onto a Formvar/Carbon 300 Mesh TEM grid (Electron Microscopy Sciences, FCF300-CU). It was dried overnight inside a Petri dish at 4 °C, which was shielded with parafilm and disinfected with 100% EtOH prior to grid preparation. Before performing the TEM analysis, the vesicle samples were covered with another 300-mesh TEM grid. The sample was then imaged using Transmission Electron Microscope (JEOL, JEM-2100), with 5 data points obtained from different TEM images in three independent experiments.

### Zeta potential measurement

For zeta potential measurements, a Zetasizer Nano ZS (Malvern Instruments Ltd., Cambridge, UK) was utilized. To measure the zeta potential of EVs and CLIPs, the vesicles were diluted in PBS that had been passed through a 0.2 µm syringe filter, to remove particles. After the dilution, 1 mL of the solution was loaded into a disposable folded capillary zeta cell (Malvern Instruments Ltd, DTS1070). In the case of EV-CLIP measurements, following the generation of the droplet emulsion using the droplet generation chip, the droplet emulsion was centrifuged 10,000 ×*g* for 5 min. After that, the upper layer was then extracted and 1 × PBS was introduced. Subsequently, pipetting was performed, and the solution was vortexed for 30 s. Afterwards, 1 mL of the solution was loaded into a disposable folded capillary zeta cell (Malvern Instruments Ltd, DTS1070). The zeta potential value was measured using the Smoluchowski approximation.

### Concentration measurement

Individual vesicles in suspension undergoing Brownian motion were monitored using Nanoparticle Tracking Analysis (NTA) by NanoSight instrument (Nanosight NS500, Malvern Panalytical). To distinguish these vesicles individually, the sample was diluted 1000× in particle-free PBS and passed through a 200-nm filter. This process ensured that the mean particle distance exceeded the diffraction limit of the microscope. Approximately 0.6 mL of the vesicle sample was introduced into the sample chamber of the NTA instrument (Malvern Analytical NanoSight N500, Malvern, UK) equipped with a 405 nm laser. To maintain a particle count between 100 and 2000, 30-s videos were generated by adjusting the camera level to 14 and the detection threshold to 7. Data analysis was conducted using NTA 3.1 software (Nanosight) in automatic mode to adjust camera focus, shutter, blur, minimum track length, minimum expected size, and maximum jump distance. The concentrations of CLIPS, EVs, and EV-CLIPs were determined based on a minimum of three independent experiments.

### EV-CLIP colocalization experiment

For the EV-CLIP colocalization study, the EVs and CLIPs were labeled with 3,3′-dioctadecyloxacarbocyanine perchlorate (DiO) dye (V22886; Thermo Fisher Scientific) and rhodamine-DHPE, respectively. For EV labeling, 100 µL of pre-concentrated EVs from the ExoDisc platform were incubated with 2 µL of 200 nM DiO dye solution for 20 min on a shaker at room temperature. After incubation, the EVs were washed twice with 1 × PBS and centrifuged at 3000 rpm. The concentrated EVs from the chamber were resuspended in 1 × PBS with a dilution factor of two. For CLIP labeling, 1 mol% of rhodamine-DHPE was incorporated into the lipid mixture. The volume was then adjusted to reach a final lipid concentration of 5 mg/mL. The Rho-CLIPs were synthesized using the aforementioned MHF method. The synthesized CLIPs were purified by washing with PBS using a 100 KDa Amicon centrifugal filter to remove the free dyes. The dye-modified CLIPs were fused with the EVs using the abovementioned microfluidic-aided fusion method. Following generation using the droplet generation chip, the droplet emulsion was centrifuged at 10,000 ×*g* for 5 min. Subsequently, the upper layer was extracted, and 1 × PBS was introduced. The solution was pipetted and vortexed for 30 s and imaged using the confocal laser-scanning microscope (Zeiss LSM 780NLO) using a ×10 Plan-Apochromat, 0.45 NA objective lens with an LU-NV laser unit 488 nm (green/Alexa Fluor 488) and 560 nm (red/Cy5). The colocalization analysis was performed using FIJI.

### NBD-PE fusion quantification assay

Fusion quantification was conducted using a previously published method (*52*). Briefly, 1 mol% of rhodamine-DHPE and NBD-PE were incorporated into the lipid mixture. The volume was then adjusted to reach a final lipid concentration of 5 mg/mL. The Rho-NBD CLIPs were synthesized using the aforementioned MHF method. The synthesized CLIPs were purified by PBS washing using the 100 KDa Amicon centrifugal filter to remove the free dyes. Dye-modified CLIPs were fused with the EVs using the abovementioned microfluidic-aided fusion method. The fluorescence signal from the fused vesicles was measured using a spectrophotometer (M2000 Pro; Tecan) with an excitation/emission wavelength of 485/535 nm, respectively. The fusion percentage was then calculated using the following formula:

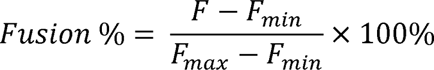

where F_max_ is the fluorescence intensity of the fused vesicles lysed with Triton-X, and F_min_ is the fluorescence intensity of EVs.

### Molecular beacon loading into the CLIPs

The miR-21, EGFR L858R, and EGFR T790M targeting molecular beacons (Macrogen, Korea) were loaded into the CLIPs by incorporating 1 µM of molecular beacons in 1 × PBS (pH 7.4) at 25 L. The CLIPs were synthesized using the MHF method with a final lipid concentration of 5 mg/mL. miR-21-targeting beacon was loaded into a batch of CLIPs, whereas EGFR L858R and T790M mutation-specific beacons were loaded simultaneously into a different batch of CLIPs. Pump flow rates were maintained at 50 and 5 µL/min for the aqueous and oil phases, respectively. The molecular beacon-loaded CLIPs were analyzed using a Nanosight instrument to determine the concentration and size and Malvern Zetasizer to determine zeta potential distribution.

### EV-derived RNA detection using the EV-CLIP method

EVs were detected using the microfluidic cascade platform, which integrates the MHF chip for CLIP generation, a droplet-based fusion of CLIPs and EVs, and droplet scanning into the detection chamber chip. Briefly, CLIPs containing a fixed concentration (1 µM) of MB (miR-21 or EGFR L858R and T790M mutations) were generated using the aforementioned method. These CLIPs were then introduced into the droplet generation chip, where fusion occurred with various concentrations of EVs (10–10^5^ particles/µL) within 30-µm diameter droplets generated in the FC-40 oil phase. Pump pressures were maintained at 2500 and 1100 mBar for oil and aqueous phases, respectively. The droplet solution was collected for 20 min in 1.5 mL brown microcentrifuge tubes and directly pumped into the detection chamber chip, in which the signal was observed under the confocal fluorescent microscope (Zeiss LSM 780NLO, ×10 Plan-Apochromat, 0.45 NA objective, LU-NV laser unit 488 nm (green/Alexa Fluor 488) and 560 nm (red/Cy5), Laser: 6.5%, Pinhole: 1.46 Airy Units, Laser gain: 750, Digital gain: 1.6, Speed: 7, Acquisition time: 7.75 s).

### Experiments with EVs spiked in plasma from healthy donors

Institutional Review Board (IRB)-approved healthy human plasma was obtained from the Red Cross (UNISTIRB-19-41-C). Briefly, the human plasma was spiked with H1975-derived EVs at different concentrations (10–10^5^ particles/µL) without prior dilution. The spiked plasma samples and MB-loaded CLIPs (miR-21 or EGFR L858R and T790M mutations) were introduced into the droplet chip for microfluidic-aided fusion. Pump pressures were maintained at 2500 and 1100 mBar for oil and aqueous phases, respectively. The droplets were collected for 20 min in microcentrifuge tubes and immediately pumped into the detection chamber chip.

### EGFR mutation detection in patient sample

All patient samples used were approved by the IRB. Eighty-three samples—10 healthy donor samples, 17 lung cancer samples without mutation (6 stage I, 3 stage II, 3 stage III, and 5 stage IV samples), and 56 lung cancer samples with EGFR mutations (18 stage I, 6 stage II, 7 stage III, and 25 stage IV samples)—were obtained from Inha University Hospital (2019-11-017), Chonnam National University Hwasun Hospital (CNUHH-2022-021), and Pusan National University Hospital (2106-044-104). The detection procedure was identical to the previously described detection method. Briefly, while the generated CLIPs were directly connected to the droplet chip, 20 µL of the samples were loaded into the droplet chip for microfluidic-aided fusion. Pump pressures were maintained at 2500 and 1100 mBar for oil and aqueous phases, respectively. The droplets were collected for 20 min in microcentrifuge tubes and pumped into the imaging chamber chip for further analysis.

### Threshold Determination for Image Analysis

The threshold for the image analysis was determined to be 56 and 70 in 8-bit 0–255 scale for miR21, L858R (green dyes) and T790M (red dyes) respectively. The threshold was calculated by mean of maximum + 3SD value of healthy donor.

### Image processing and analysis

For imaging, 6 µL of the droplet solution was loaded into the 30 µm chamber chip (10001447; Microfluidic ChipShop), and the droplets were observed in the confocal laser-scanning microscope (Zeiss LSM 780NLO) using a ×10 Plan-Apochromat, 0.45 NA objective lens with an LU-NV laser unit 488 nm (green/Alexa Fluor 488) and 560 nm (red/Cy5). The sample was imaged with the parameters of 6.5% laser power, pinhole size of 1.46 airy units, laser gain of 750, digital gain of 1.6, Speed of 7 with acquisition time of 7.75 s with 10 frames taken for auto stacking mode.

After imaging, all images for droplet analysis were processed using FIJI. The code describes the processing steps for image quantification.

#### Image breakdown (splitting the image)

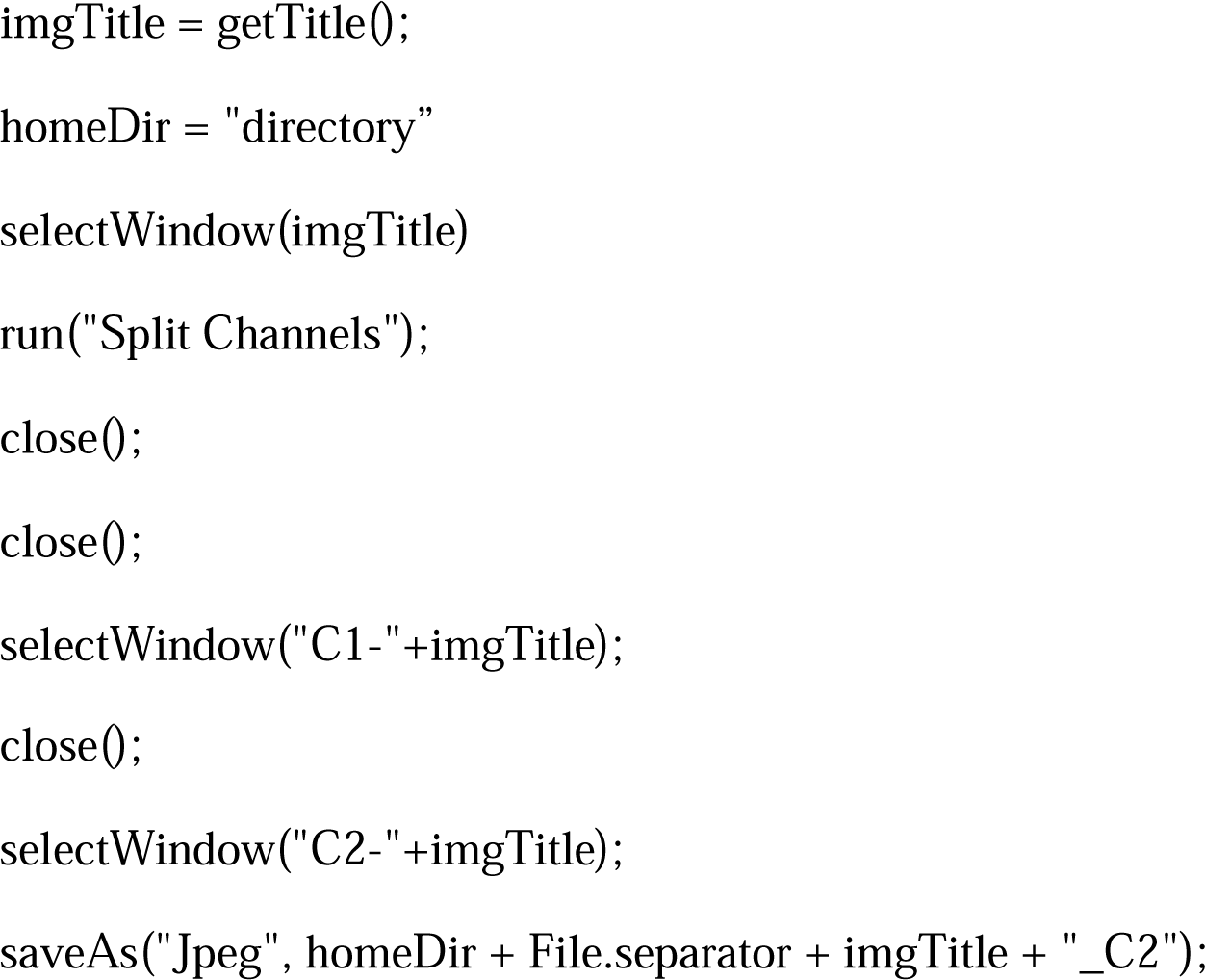

#### Brightfield processing

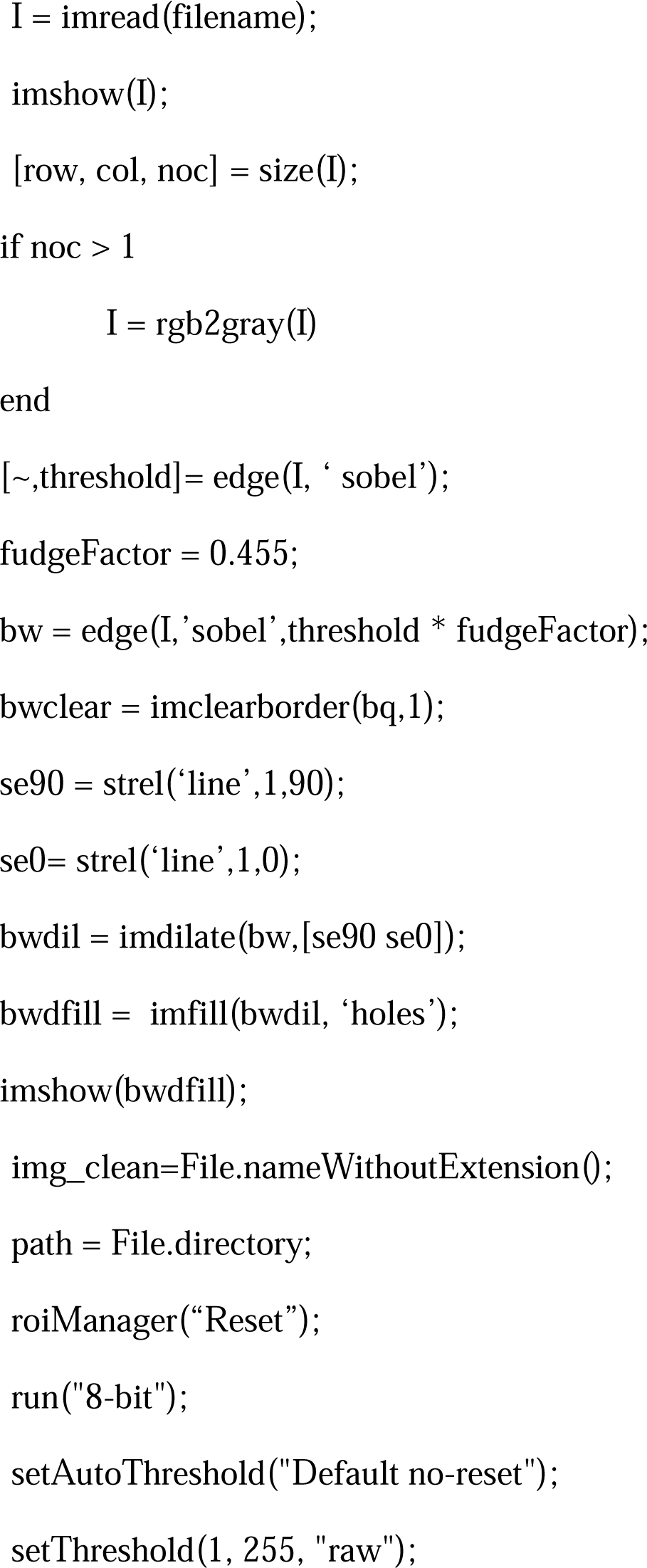
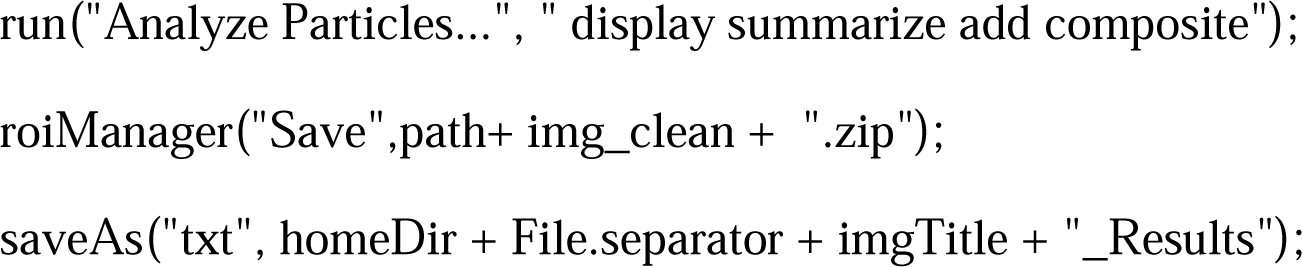

#### Fluorescent processing

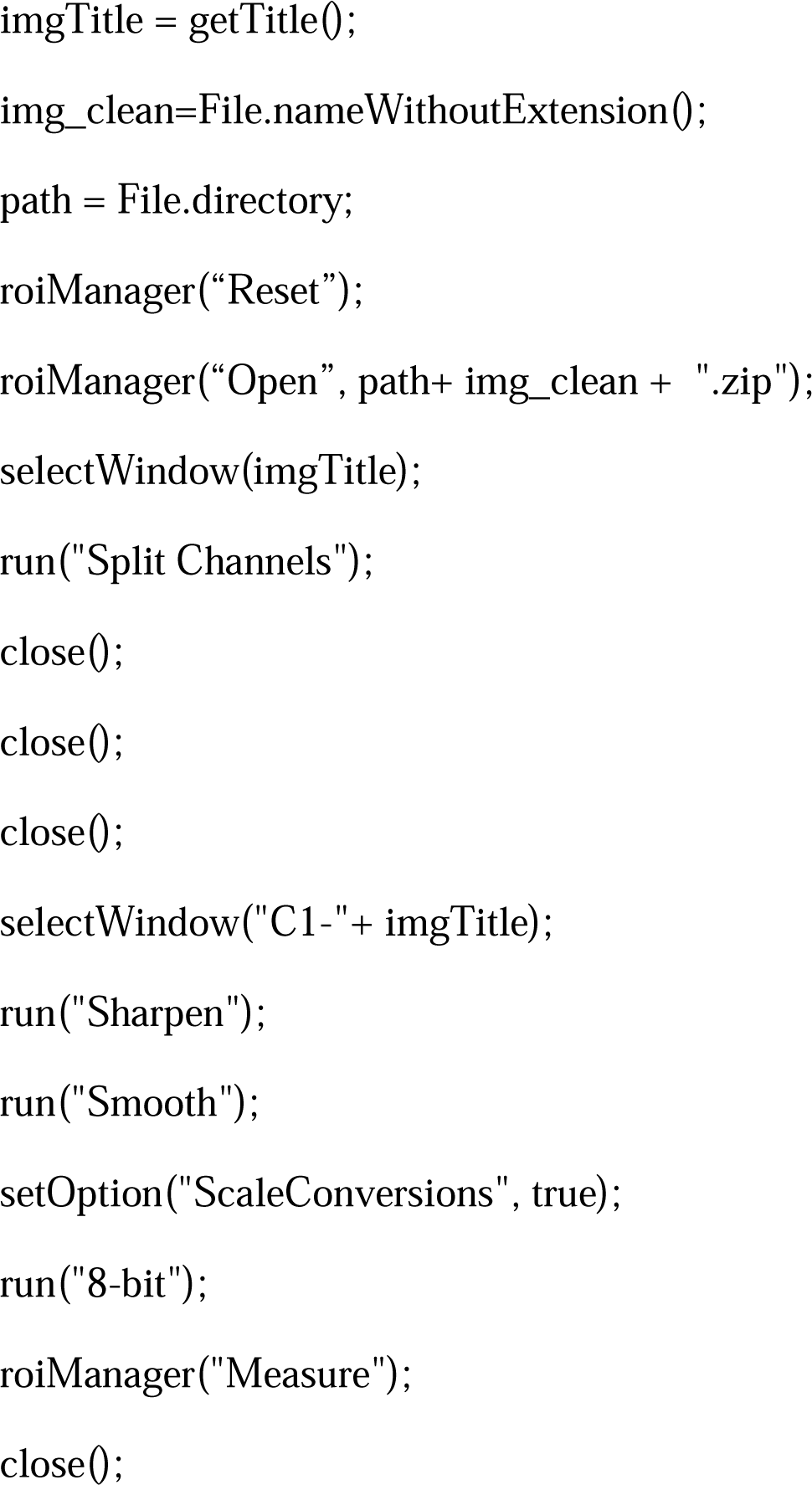

### Droplet digital PCR (ddPCR) of EV-spiked human plasma

ddPCR was performed to compare our platform with a pre-existing platform. Briefly, healthy donor human plasma, obtained from the Red Cross (UNISTIRB-19-41-C), was spiked with H1975 EVs at concentrations varying from 10 to 10^5^ particles/µL. EVs from the sample were separated using the ExoRNeasy midi kit (Qiagen). After EV separation, RNA was extracted, and cDNA was synthesized using the SuperScript VILO cDNA synthesis kit (Thermo Fisher Scientific). ddPCR samples were prepared with 10 μL of ddPCR Super Master Mix (No dUTP), 5 μL of cDNA template, 1 μL of each primer/probe mix, and 3 μL of nuclease-free water, with a total reaction volume of 20 µL. The L858R and T790M primer/probe and mastermix were purchased from Bio-Rad (#1863104 and #1863103; Hercules, CA, USA). Droplets were generated using a QX200 droplet generator (Bio-Rad), using 70 μL of oil as the probe. PCR amplification was executed using the Mastercycler Pro S (Eppendorf, Hamburg, Germany) with the configuration as follows: 95 °C for 10 min; 40 cycles of 94 °C for 30 s, 58 °C for 1 min, and 98 °C for 10 min, followed by a hold at 4 °C.

### Statistical analysis

Experiments were independently conducted at least three times, with the number of replicates specified in each figure legend. Two-group comparisons were performed using a two-tailed unpaired Student’s *t*-test, and three- or more group comparisons were conducted using one-way ANOVA. Statistical analyses were performed using GraphPad Prism 10.0.0. All data represent mean ± standard deviation (SD).

## SUPPORTING INFORMATION

Materials and Methods

Fig S1 to S29

Tables S1 to S5

Movies S1 to S2

References (*51-67*)

## ACKNOWLEDGMENT

This work was supported by the Center for Soft and Living Matter, Institute for Basic Science (IBS-R020-D1) by the Korean Government and the BK21 program through the National Research Foundation (NRF) funded by the Ministry of Education of Korea.

## AUTHOR CONTRIBUTIONS

All authors contributed to writing the manuscript. All authors have approved the final version of the manuscript.

## DATA AND MATERIALS AVAILABILITY

All data associated with this study are present in the paper or the Supplementary Materials.

