## Supplementary Materials for "Digital Profiling of Tumor Extracellular Vesicle-associated RNAs Directly from Unprocessed Blood Plasma"

Materials and Methods

Fig. S1 to S29

Tables S1 to S5

Movies S1 to S2

References (*51-67*)

**Materials and Methods**

**Reagent & materials**

We used a 0.2 µm syringe filter (S6534; Minisart® NML Syringe Filters), 1×PBS (pH 7.4, Gibco), 50 g of 2 wt.% 008-FluoroSurfactant in FC40 (0089-fluorosurfactant-2wtF-50G; RAN Biotechnologies), antibiotic/antimycotic (15240062; Gibco) EV-depleted FBS, Exodisc^TM^-C (LabSpinner), molecular beacons (Oligo, Macrogen), NBD-PE (N-(7-nitrobenz-2-oxa-1,3-diazol-4-yl)-1,2-dihexadecanoyl-sn-glycero-3 phosphoethanolamine, triethylammonium salt, Thermofisher, N360), and rhodamine-DHPE (Lissamine rhodamine B 1,2 dihexadecanoyl-sn-glycero-3-phosphoethanolamine, triethylammonium salt, L1392; Thermo Fisher Scientific).

Additionally, we used Droplet Chip (3200529; Dolomite Microfluidics), 150 µm microfluidic hydrodynamic focusing (MHF) chip 3D (3200834; Dolomite Microfluidic), 30 µm chamber chip (10001447; Microfluidic ChipShop), 96-well plate (3364; Corning), confocal laser-scanning microscope (Zeiss LSM 780NLO, Zeiss), Malvern Zetasizer (Nano ZS), NanoSight instrument (Nanosight NS500, Malvern Panalytical), M2000 Pro Plate Reader (Tecan), and transmission electron microscope (JEM-2100; JEOL).

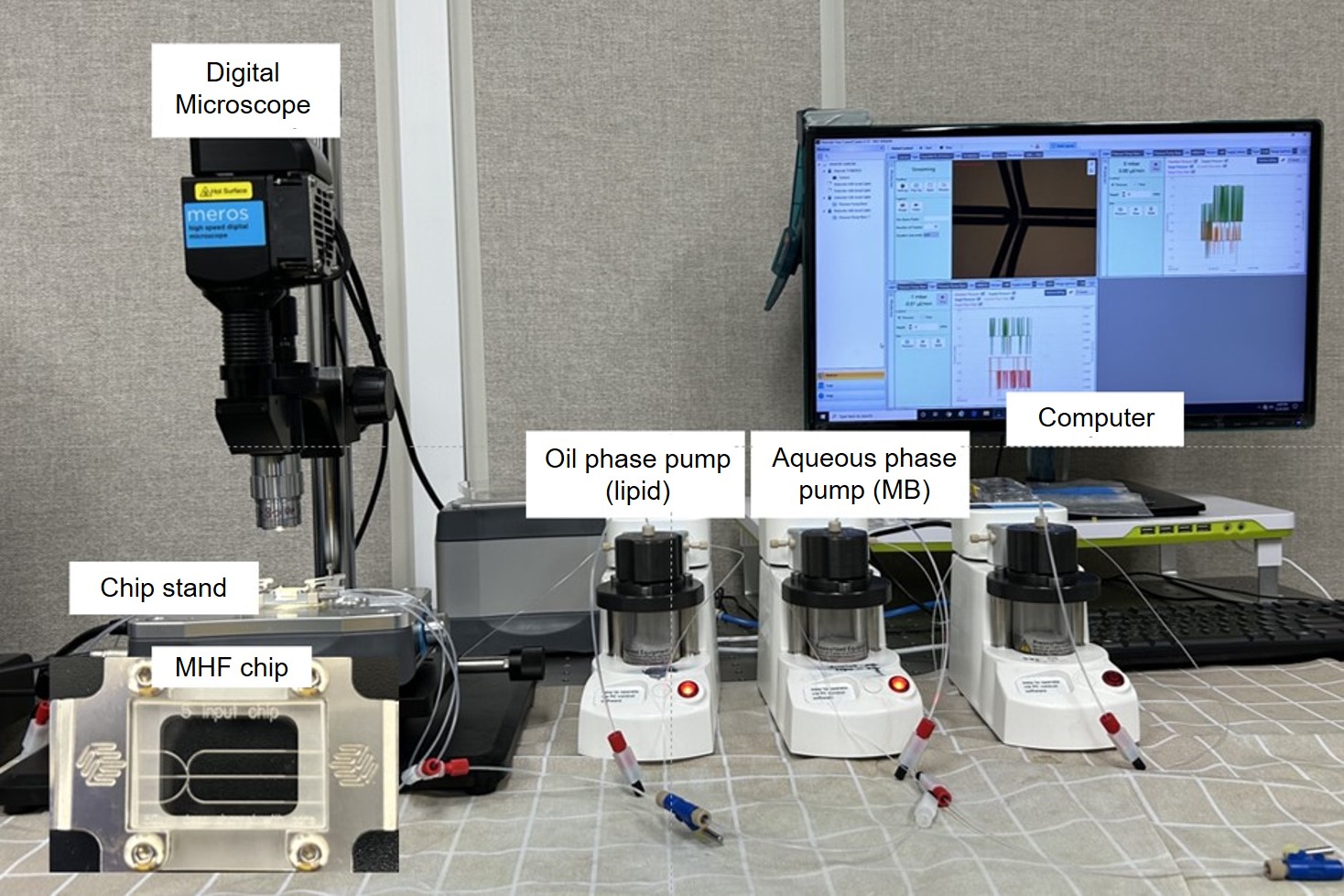

**Figure S1. Microfluidic hydrodynamic focusing (MHF)-based charged-liposome (CLIP) generation experimental setup.** The setup consists of a digital microscope, chip stand, oil phase pump, aqueous phase pump, computer, and microfluidic chip. Briefly, the lipid dissolved in ethanol (EtOH) is loaded inside the oil phase pump, and the aqueous pump is loaded with PBS. The oil phase pump pushes the lipid into the left side of the chip, and the aqueous phase pump pushes the PBS into the right side of the chip. With proper flow control, the oil layer will be sandwiched between the two aqueous layers, forming CLIPs inside the microfluidic chip. The flow rates of each pump are maintained at 50 µL/min and 5 µL/min for aqueous and oil phases, respectively.

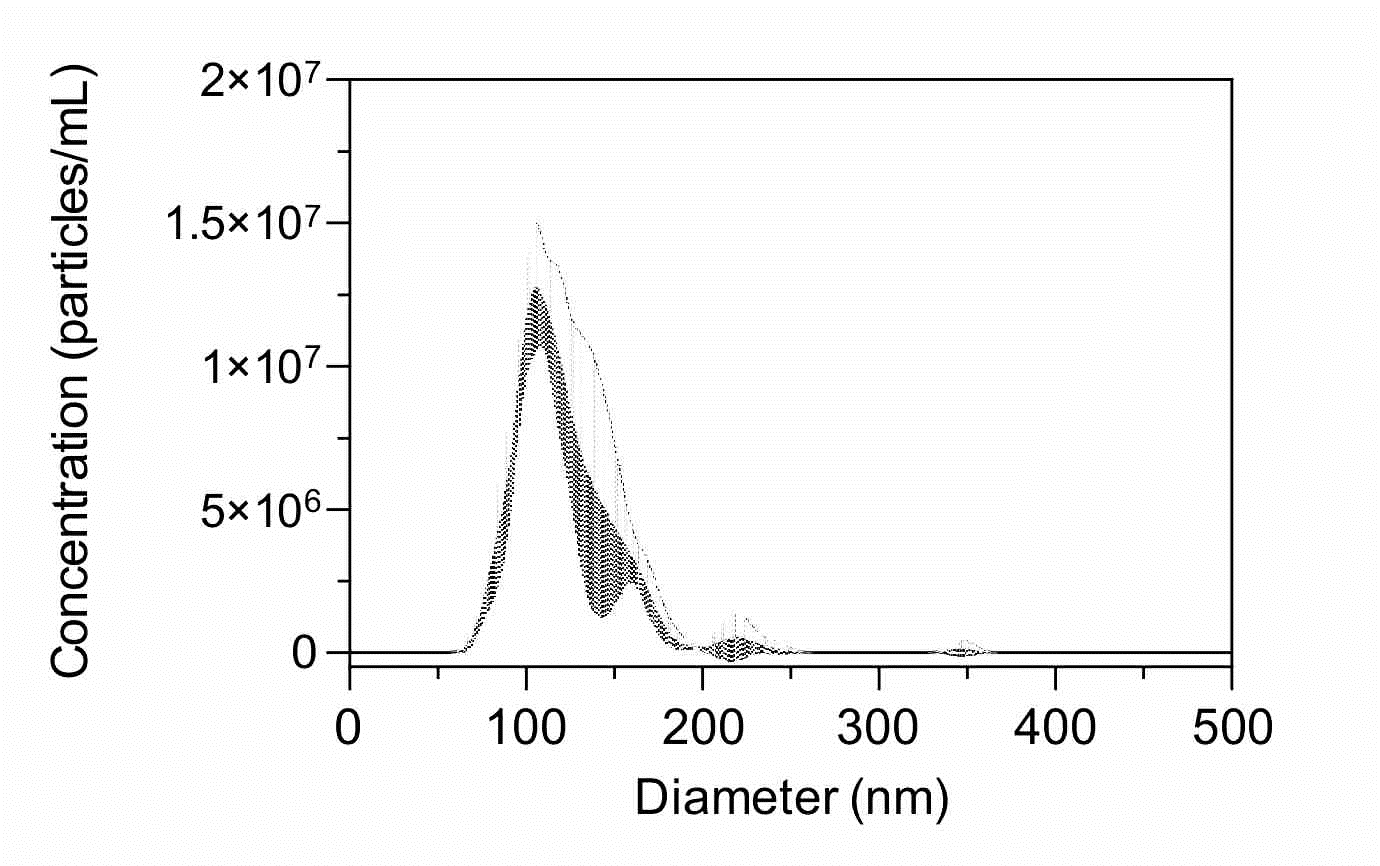

**Figure S2. Size distribution of CLIPs measured using nanoparticle tracking analysis (NTA).** Synthesized CLIPs washed using the Amicon filter were diluted in PBS and measured using NTA. The CLIPs showed a single-peak distribution curve, indicating a homogeneous CLIP population.

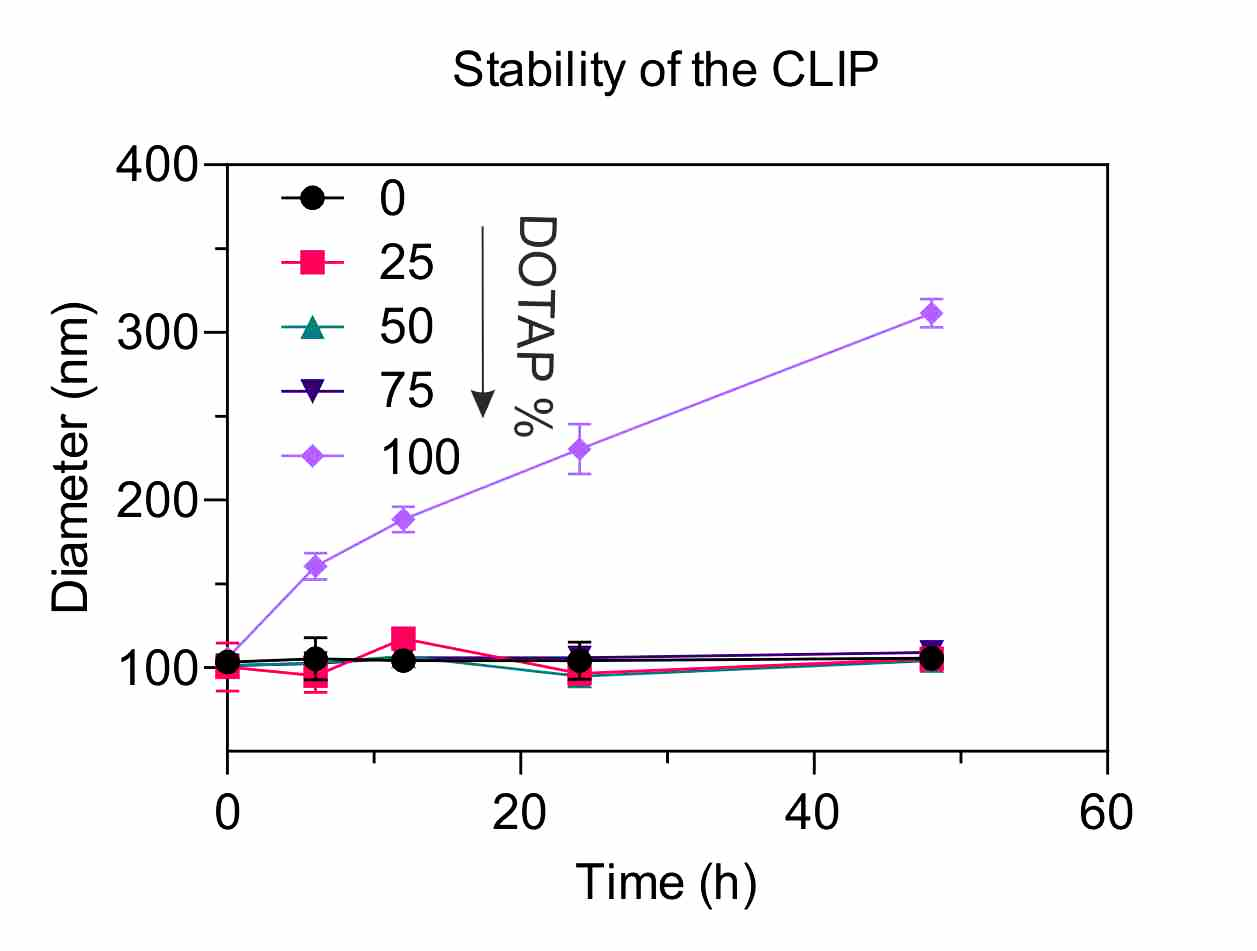

**Figure S3. CLIP stability with various DOTAP percentages over time.** Time-dependent size measurement was conducted to evaluate the size change of CLIPs containing various DOTAP percentages over 48 h. CLIP size with 0, 25, 50, and 75 mol% DOTAP remained stable over time, but that with 100 mol% DOTAP constantly increased up to 300 nm after 48 h (6 h: 160.53 ± 8.15 nm, 12 h: 188.27 ± 7.58 nm, 24 h: 230.37 ± 14.96 nm, 48 h: 311.50 ± 8.51 nm).

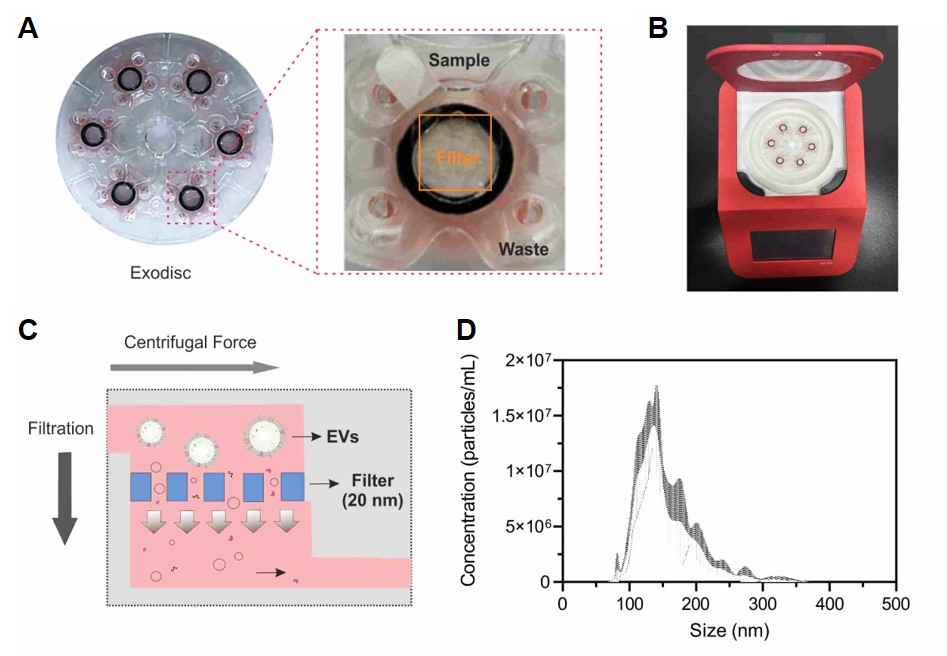

**Figure S4. EV isolation from cell culture supernatant using a lab-on-a-disc.** (**A**) Exodisc Platform, a centrifugal disk equipped with anodized aluminum oxide filters with a pore diameter of 20 nm (Exodisc^TM^-C, LabSpinner). The supernatant was passed through the filter by centrifugation at 3000 rpm (approximately 500 ×*g*), and 100 µL of the concentrated EVs from the collection chamber was resuspended in 1 × PBS with a dilution factor of two. (**B**) Tabletop centrifuge platform for Exodisc operation. (**C**) Exodisc working principle: Centrifugal force pushes the cell culture supernatant from the sample chamber through the filter chamber, with particles smaller than 20 nm being filtered out in the waste chamber. (**D**) EVs were characterized using nanoparticle tracking analysis (NTA).

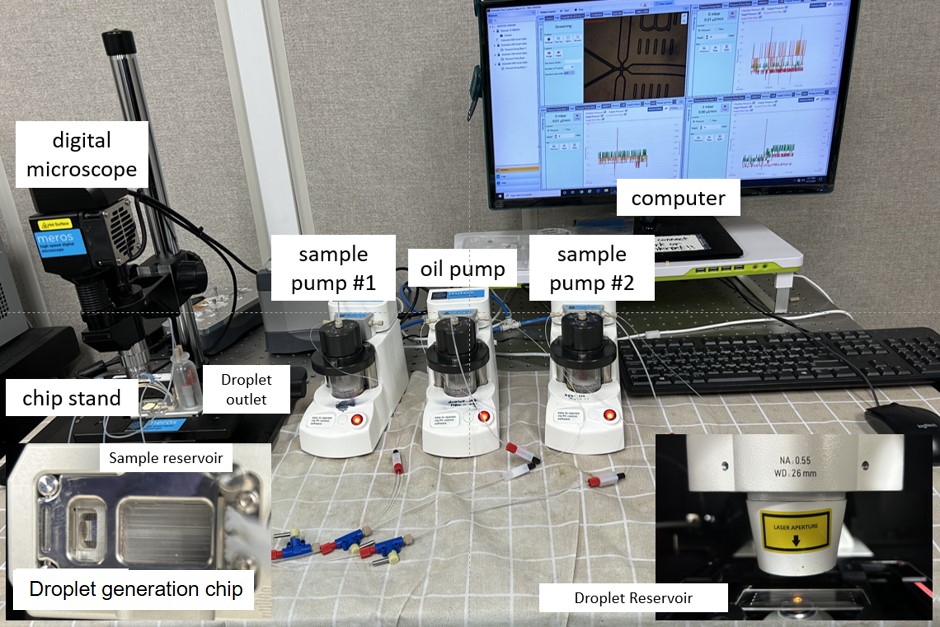

**Figure S5. Droplet generation experimental setup.** The setup comprised a digital microscope, chip stand, oil pump, two sample pumps, sample reservoir, droplet outlet, droplet chip, computer, and droplet reservoir chip. Briefly, the EVs and CLIPs are separately loaded in the two sample pumps, and the FC-40 surfactant is loaded in the oil pump and into the sample reservoir. The oil phase pump pushes the FC-40 into the vertical sides of the T junction, and the sample pumps push both EVs and CLIPs into the sample channels at 35 µL/min and 1.5 µL/min for oil and aqueous phases, respectively. The EVs and CLIPs meet at the T junction and form water-in-oil droplets collected in the droplet outlet. The droplets are imaged for analysis using the droplet reservoir chip under the fluorescence microscope.

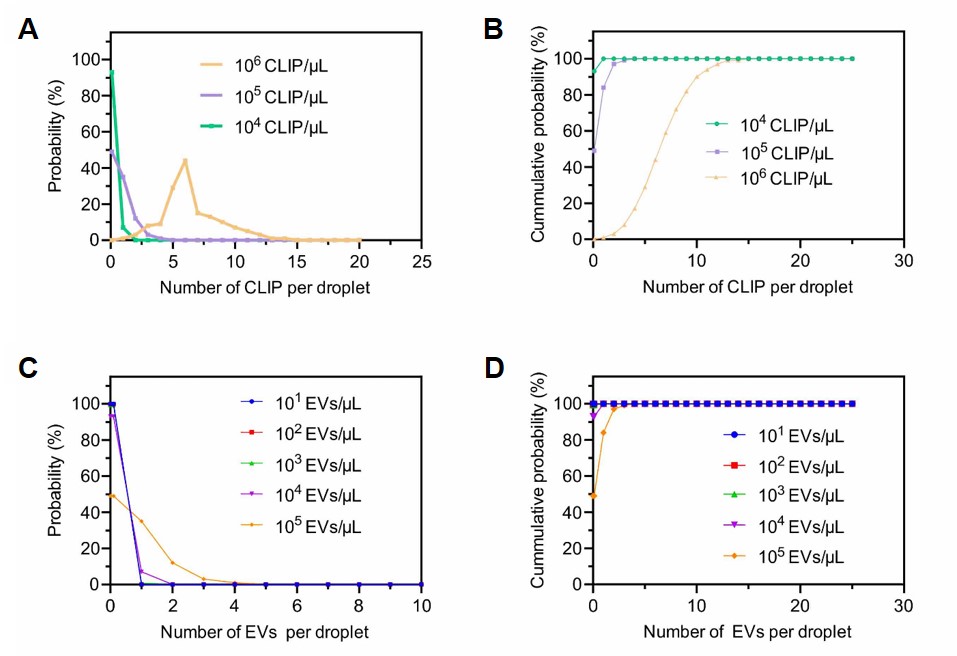

**Figure S6. CLIP and EV distribution in each droplet.** (**A**) Poisson distribution, (𝑘, 𝜆) showing the probability of droplets containing k number of CLIPs depending upon the input concentration, where λ is the average number of CLIPs per droplet volume (droplet volume = 10.77 pL; λ = 0.07, 0.71, and 7.07 for the concentrations of 10^4^, 10^5^, and 10^6^ particles µL^−1^ respectively). (**B**) Poisson cumulative distribution shows that more than 90% of droplets contain 0.1, 1–3, and 3–10 CLIPs at the optimum experimental concentration of 10^4^, 10^5^, and 10^6^ particles µL^−1^, respectively. (**C**) Poisson distribution, (𝑘, 𝜆) showing the probability of droplets containing k number of EVs depending upon the input concentration, where λ is the average number of EVs per droplet volume (λ = 0.00007, 0.00071, 0.01, 0.07, and 0.71 for the concentration of 10^1^, 10^2^, 10^3^, 10^4^, and 10^5^ particles µL^−1^, respectively). (**D**) Poisson cumulative distribution shows that more than 95% of droplets contain 0.00001–0.1 EVs at the optimum experimental concentration of 10^1^, 10^2^, and 10^3^ and 0.1–1 and 0.1–3 EVs for 10^4^ and 10^5^ particles µL^−1^.

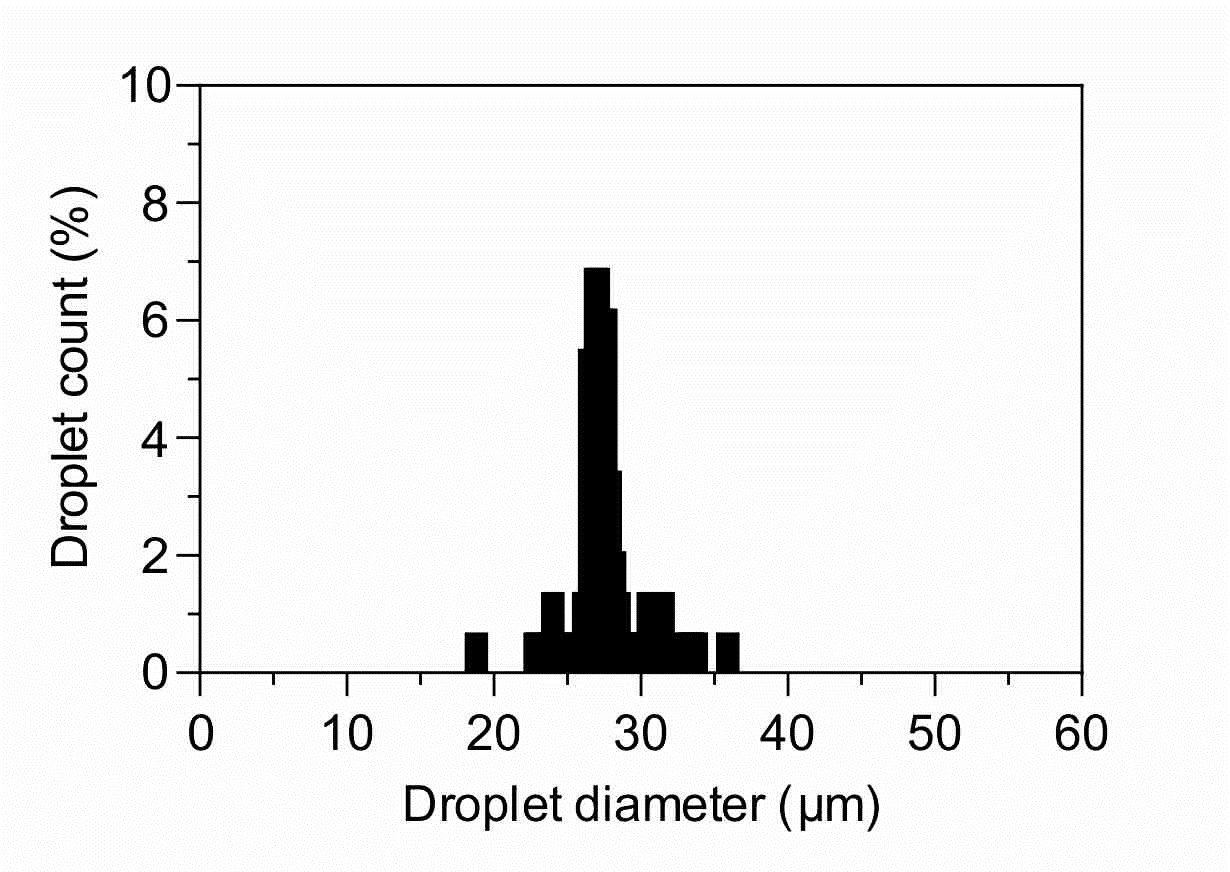

**Figure S7. Droplet size distribution graph.** The droplet was analyzed using droplet monitor software through the video taken during the droplet generation. The average size of the measured droplets was 27.40 ± 2.06 µm.

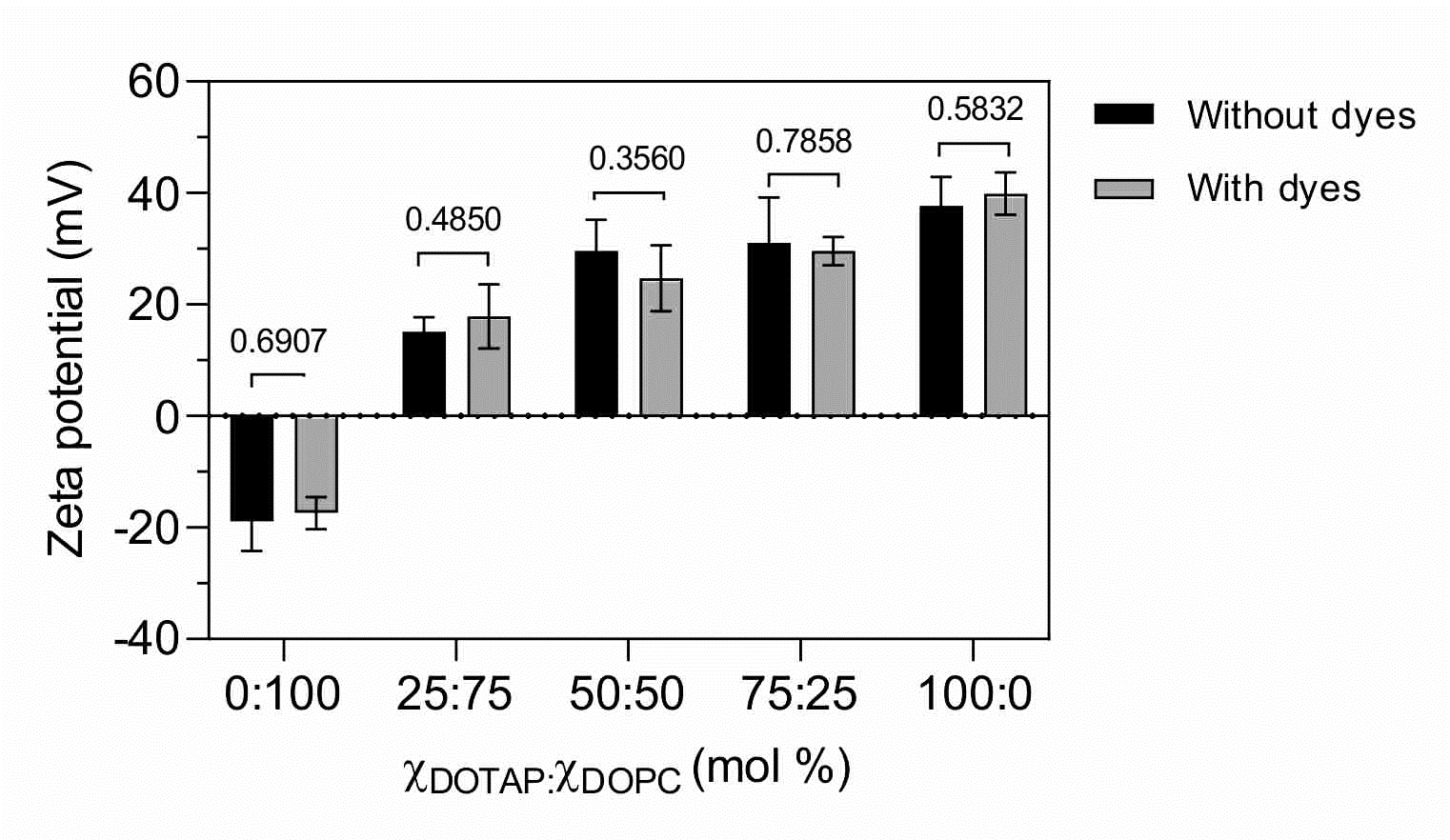

**Figure S8. Effect of Rho-NBD addition to the CLIPs on zeta potential.** Change in the zeta potential of CLIPs with various DOTAP percentages (0, 25, 75, and 100 mol%). The zeta potentials of various CLIPs without dyes are as follows: 0 mol% DOTAP, –18.9 ± 5.35; 25 mol% DOTAP, 15.10 ± 2.60; 50 mol% DOTAP, 29.60 ± 5.62; 75 mol% DOTAP, 31.00 ± 8.20; 100 mol% DOTAP, 37.70 ± 5.14. The zeta potentials after the addition of dyes are as follows: 0 mol% DOTAP, –17.40 ± 2.87; 25 mol% DOTAP, 17.90 ± 5.75; 50 mol% DOTAP, 24.70 ± 5.89; 75 mol% DOTAP, 29.60 ± 2.52; 100 mol% DOTAP, 39.90 ± 3.80. Data represent mean ± SD, n = 3. All statistical analyses were conducted using a two-tailed unpaired Student’s *t*-test.

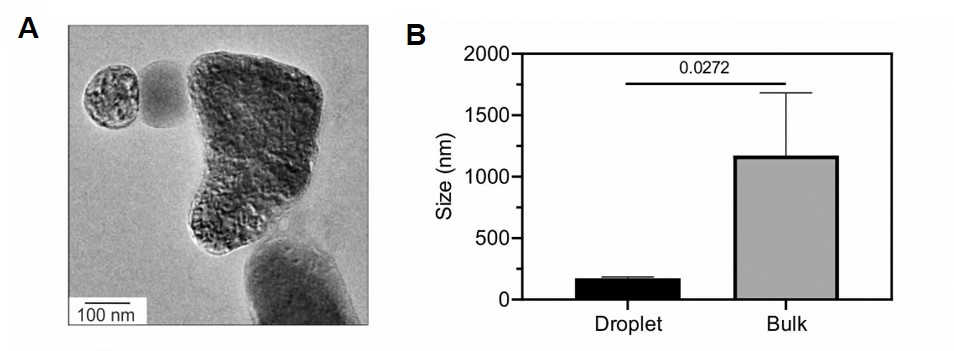

**Figure S9. TEM image and size comparison of the aggregates formed during EV-CLIP fusion in bulk scale.** (**A**) The TEM image (top) shows aggregates formed during bulk EV-CLIP fusion. Scale bar: 100 nm. (**B**) Comparison of fused vesicle size at droplet and bulk scales. The average size of fused vesicles in the droplet is 174.1 ± 10.28 nm, and that in bulk is 1173.17 ± 508.58 nm. Each data point represents mean ± SD, n = 3. Statistical analysis was conducted using a two-tailed unpaired Student’s *t*-test.

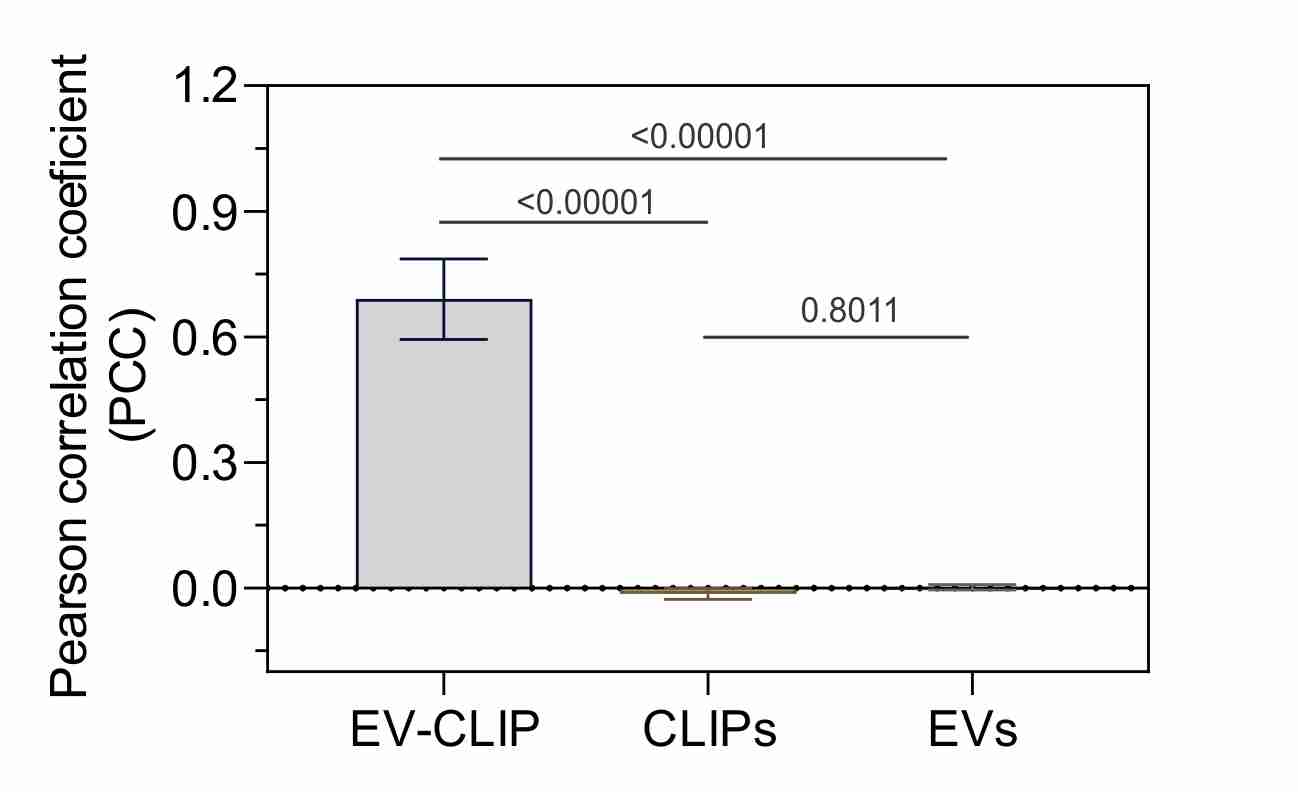

**Figure S10. Quantification of EV-CLIP vesicles, CLIPs, and EVs at different fusion ratios.** Pearson colocalization coefficient (PCC) values for three different populations. PCC values were measured from 10 different vesicle images and calculated using the JaCoP algorithm in ImageJ. The PCC was 0.690 ± 0.096, –0.0139 ± 0.01264, and 0.0021 ± 0.0066 for fused vesicles, CLIPs, and EVs, respectively. All statistical analyses were conducted using one-way ANOVA using Tukey’s multiple comparison test.

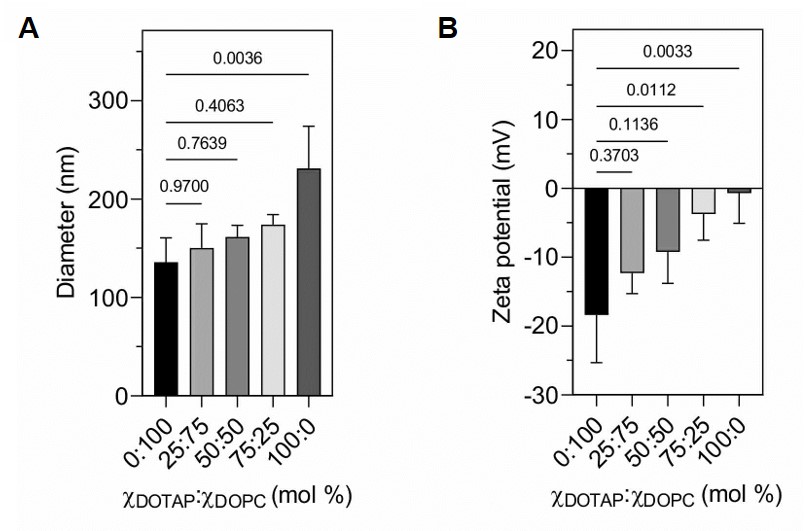

**Figure S11. Elevated size and zeta potential of the fused vesicles according to the DOTAP percentage.** (**A**) Size increase of the fused vesicles based on DOTAP percentage. (0 mol%, 135.9 ± 24.7 nm; 25 mol%, 150.5 ± 24.3 nm; 50 mol%, 161.6 ± 11.9 nm; 75 mol%, 174.1 ± 10.3 nm; 100 mol%, 231.3 ± 42.8 nm). Each data point represents mean ± SD, n = 3. All statistical analyses were conducted using one-way ANOVA. (**B**) Change in surface charge of the fused vesicles based on DOTAP percentage (0 mol%, –18.4 ± 6.9, 25 mol%, –12.3 ± 3.0, 50 mol%, –9.24 ± 4.55; 75 mol%, –3.74 ± 3.80; 100 mol%, –0.71 ± 4.39). Each data point represents mean ± SD, n = 3. All statistical analyses were conducted using one-way ANOVA using Dunnett’s multiple comparison test.

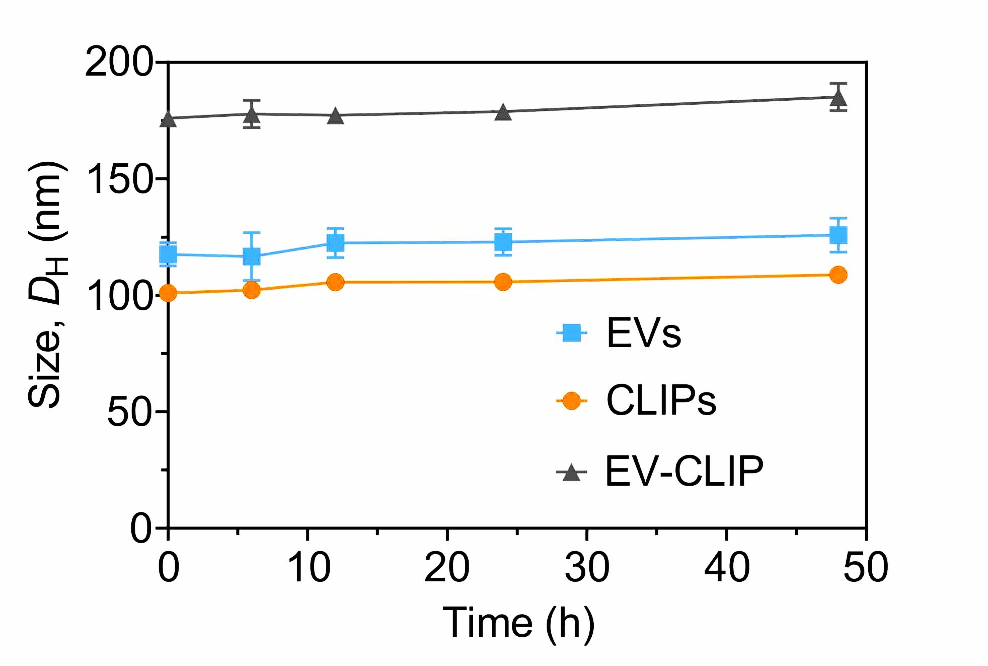

**Figure S12. Vesicle stability test after prolonged storage.** Time-dependent size measurement of the EV-CLIP, EVs, and CLIPs using the DLS. Each data point represents mean ± SD, n = 3 independent experiments.

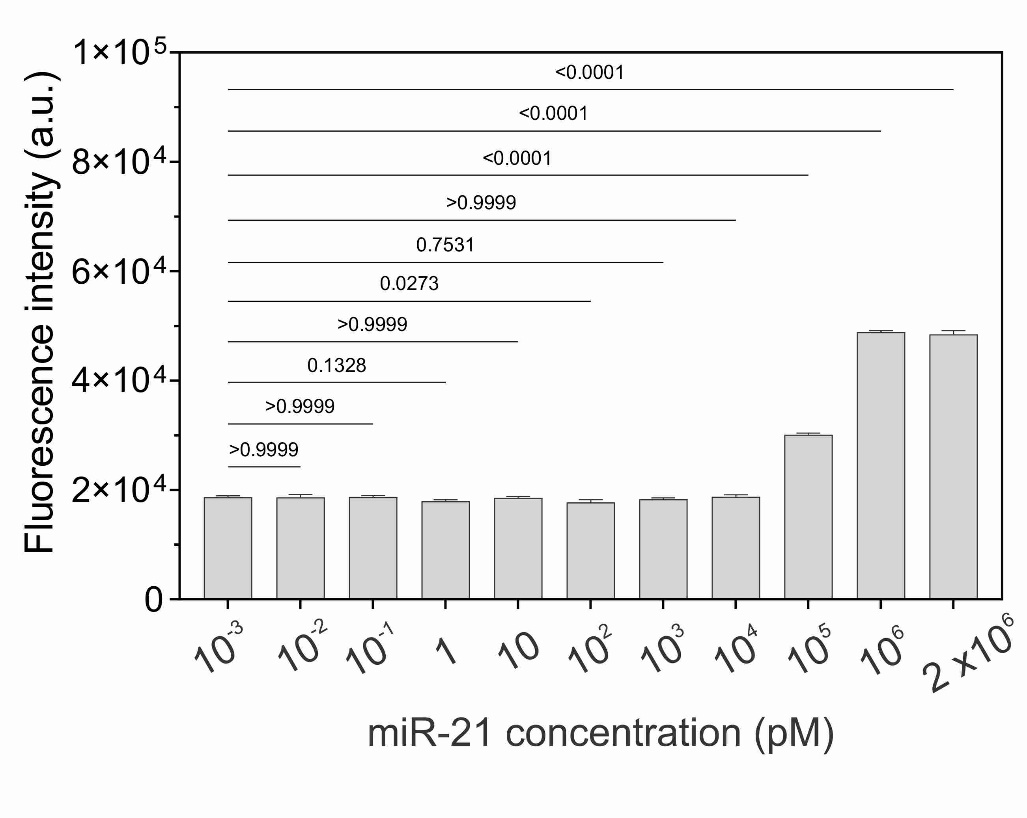

**Figure S13. Loading of the miR-21 molecular beacon into CLIPs.** Different concentrations of molecular beacons were tested, with 1 µM demonstrating the optimum signal compared to other concentrations. All statistical analyses were conducted using one-way ANOVA using Dunnett’s multiple comparison test. Each data point represents mean ± SD, n = 3 independent experiments.

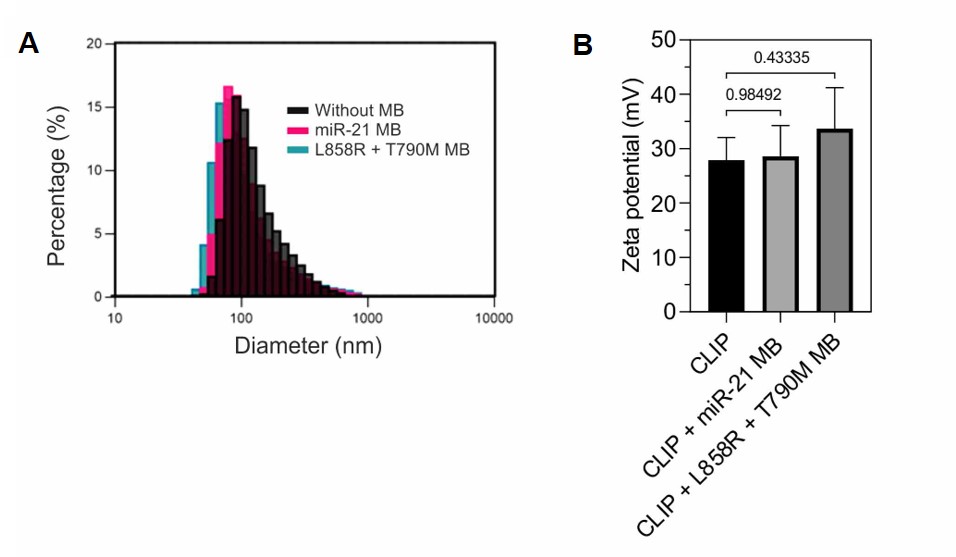

**Figure S14. Effect of MB insertion into the CLIPs in terms of size and zeta potential.** (**A**) Size distribution of three different CLIP populations: without molecular beacon (MB), with miR-21-detecting MB, and EGFR mutation-detecting MB. (**B**) Zeta potential of three different CLIP populations: without MB, 27.90 ± 4.20 mV; with miR-21-detecting MB, 28.60 ± 5.70 mV; and EGFR mutation-detecting MB, 33.70 ± 7.50 mV. Each data point represents mean ± SD, n = 3. All statistical analyses were conducted using one-way ANOVA using Dunnett’s multiple comparison test.

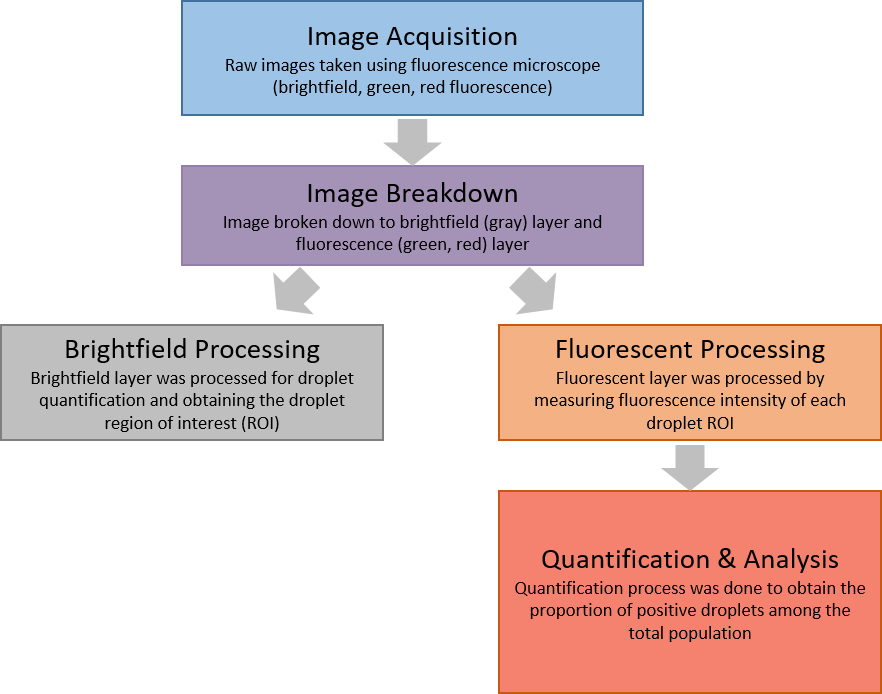

**Figure S15.** **Droplet image processing flowchart.** The flowchart outlines the sequential processing steps for droplet image analysis, beginning with image acquisition and progressing through processing, quantification, and analysis.

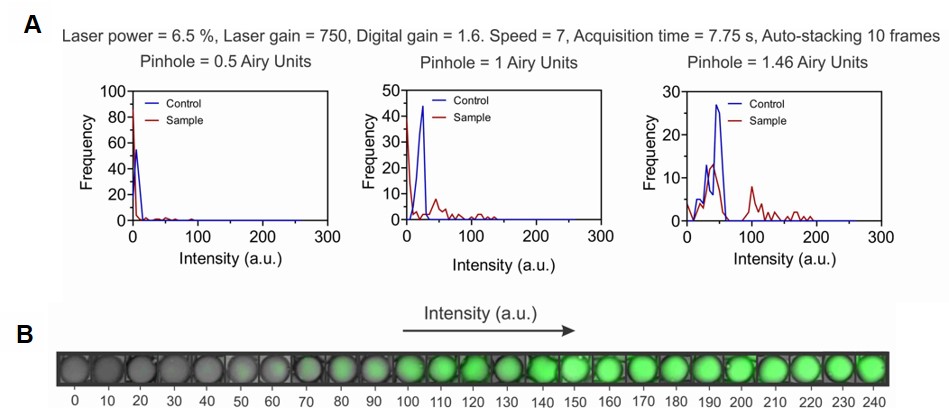

**Figure S16.** **Imaging Condition for EV-CLIP** (**A**) Imaging was done using the conditions specified. The pinhole size was increased to enhance the signal coming from the EV-CLIP. (**B**) Visualization of droplets having different intensity varying from 0 to 255.

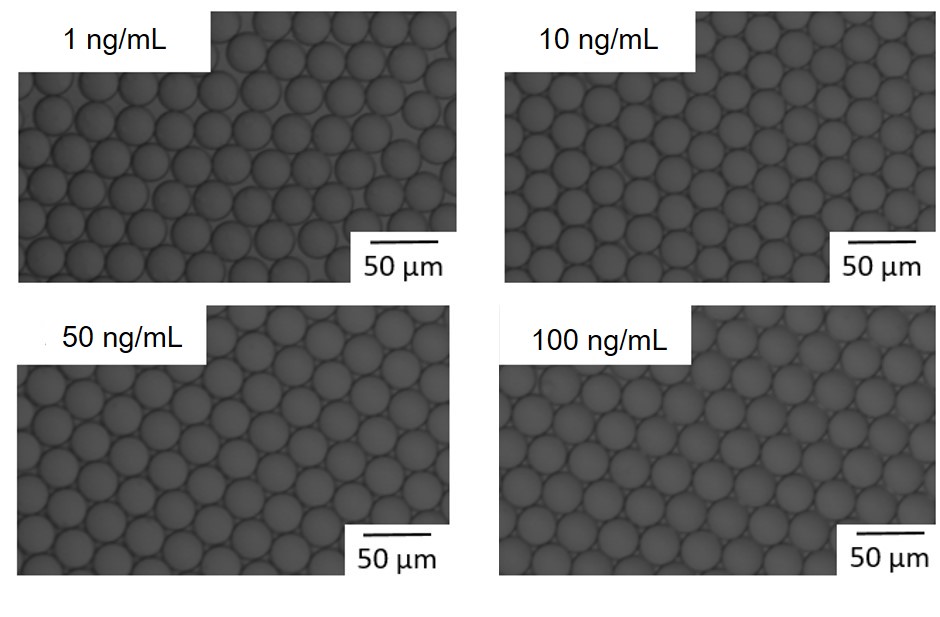

**Figure S17.** **Free miR-21 experiment under different conditions.** miR-21 concentration varied from 1 to 100 ng/mL. No signal could be detected, indicating that the fluorescent signal comes from the miR-21 inside the EVs.

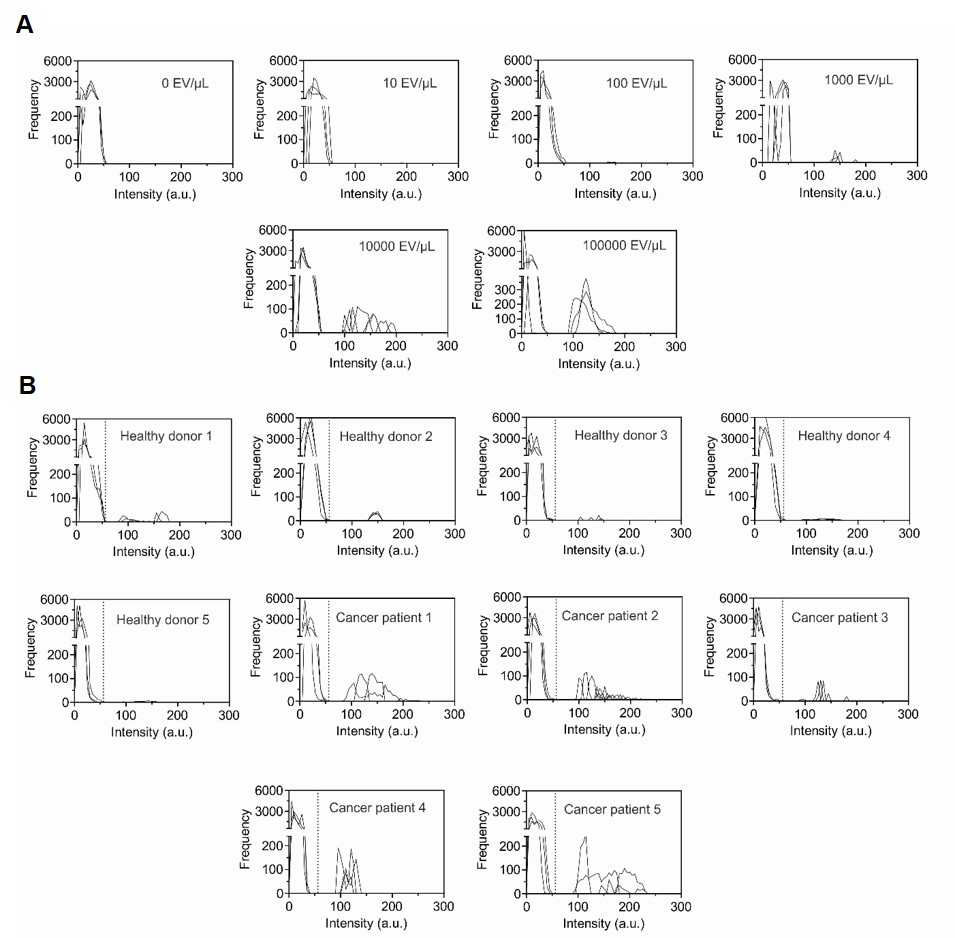

**Figure S18. Histogram of EV-CLIP Droplets for miR-21 Detection** (**A**) Histogram for intensity profile of EV-CLIP droplets having various concentrations of H1975 EVs in PBS for the detection of miR-21. The peak increases as the input EV concentration increases from 10 EVs/μL to 100,000 EVs/μL (**B**) intensity profile of EV-CLIP droplets from 5 healthy donors and 5 patients with lung cancer for the detection of miR-21. Patients with lung cancers exhibit noticeably bigger peaks compared to healthy donors, indicating that there are more miR-21 containing EVs.

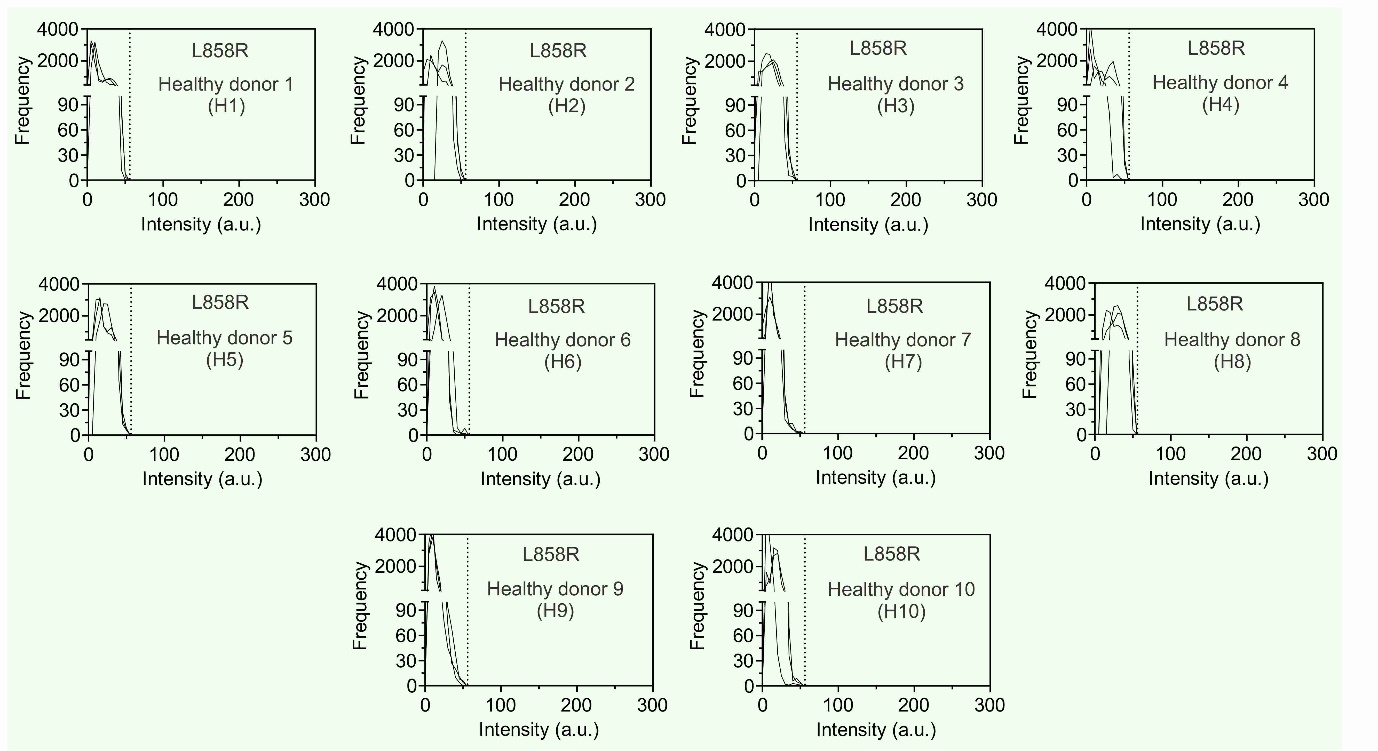

**Figure S19. Histogram of EV-CLIP droplets for L858R detection from healthy donor.** Histogram for intensity profile of L858R EV-CLIP droplets from 10 healthy donors.

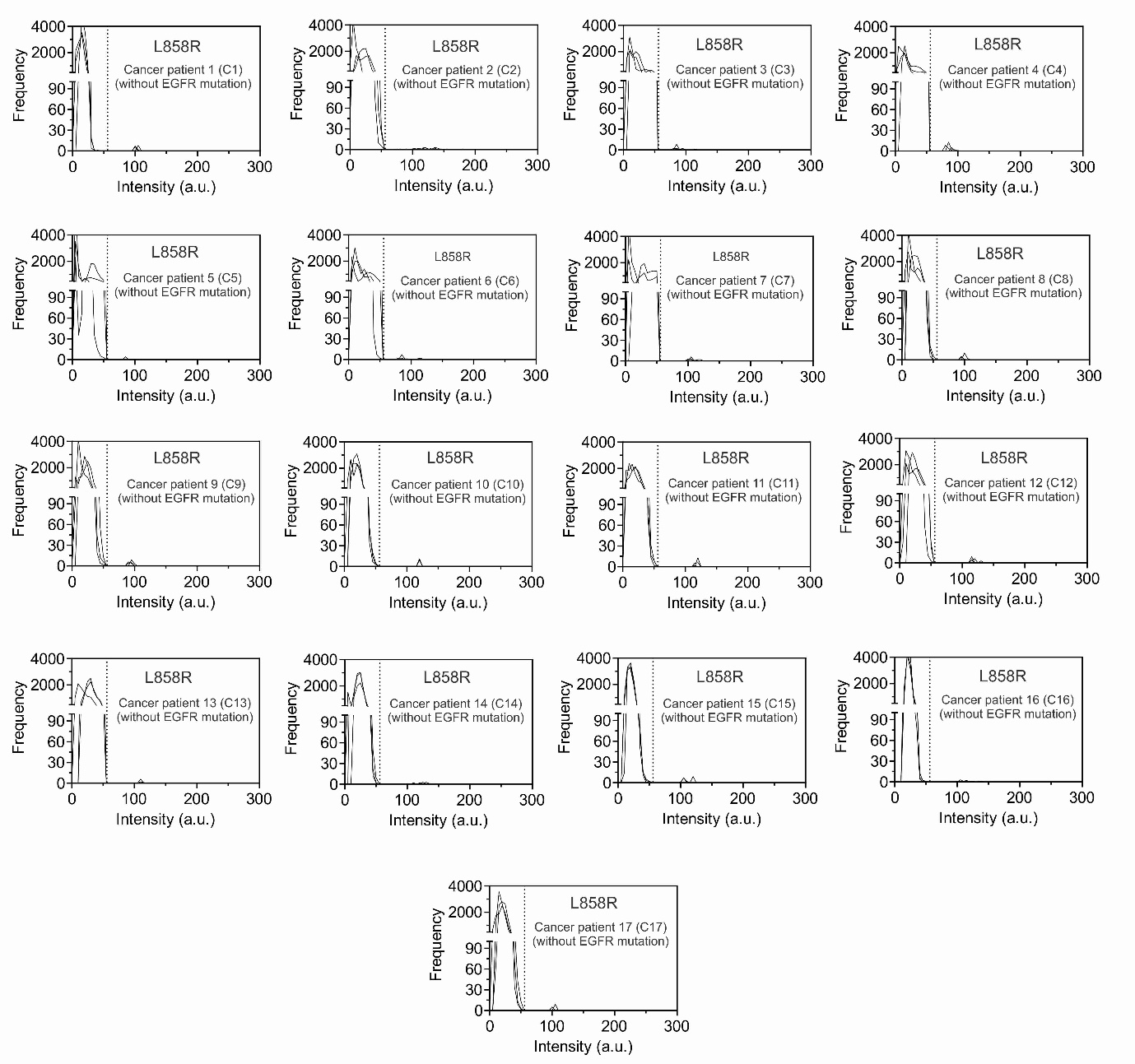

**Figure S20. Histogram of EV-CLIP droplets for L858R detection from clinical samples without EGFR mutation.** Histogram for intensity profile of L858R EV-CLIP droplets from 17 patients with lung cancer without EGFR mutations.

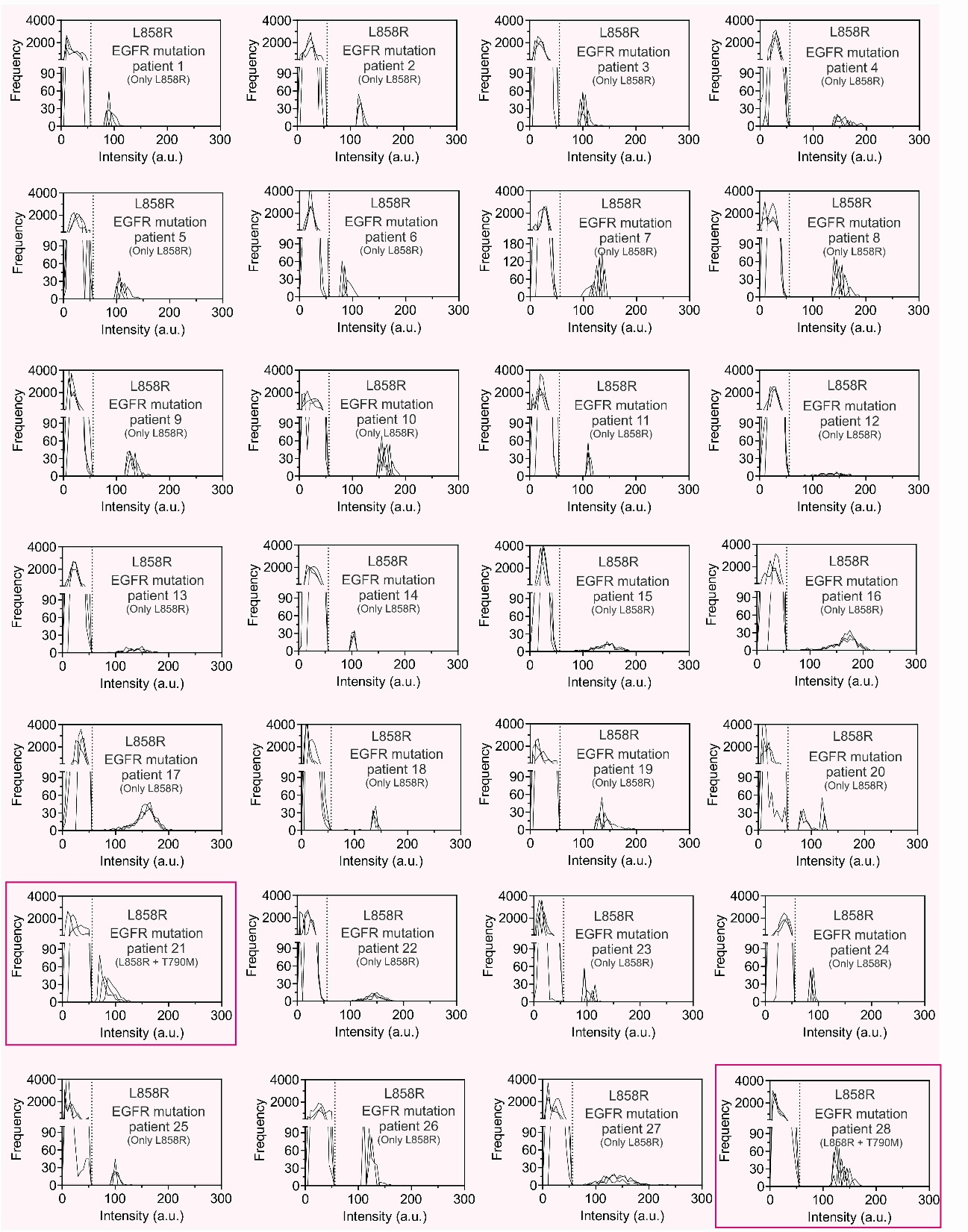

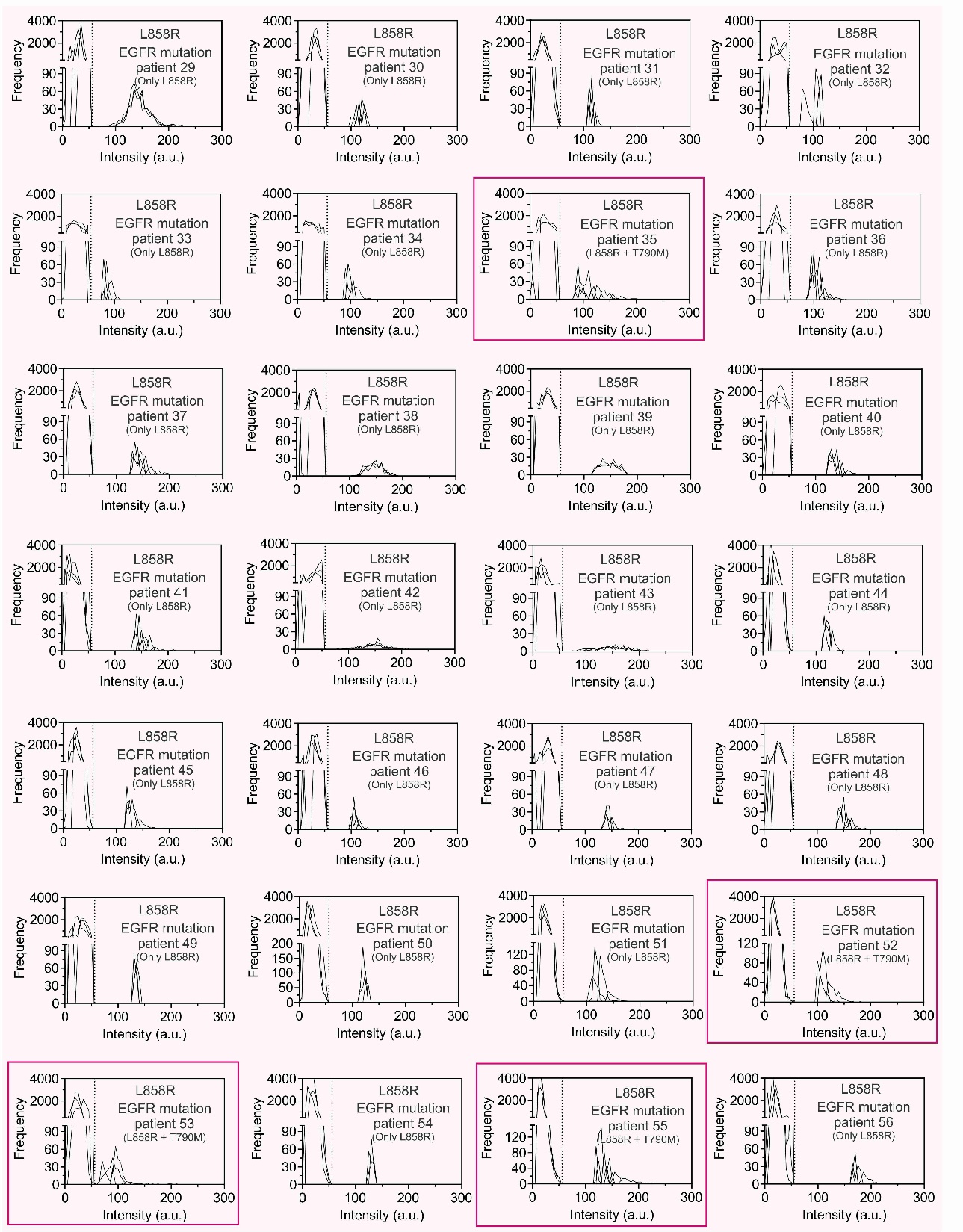

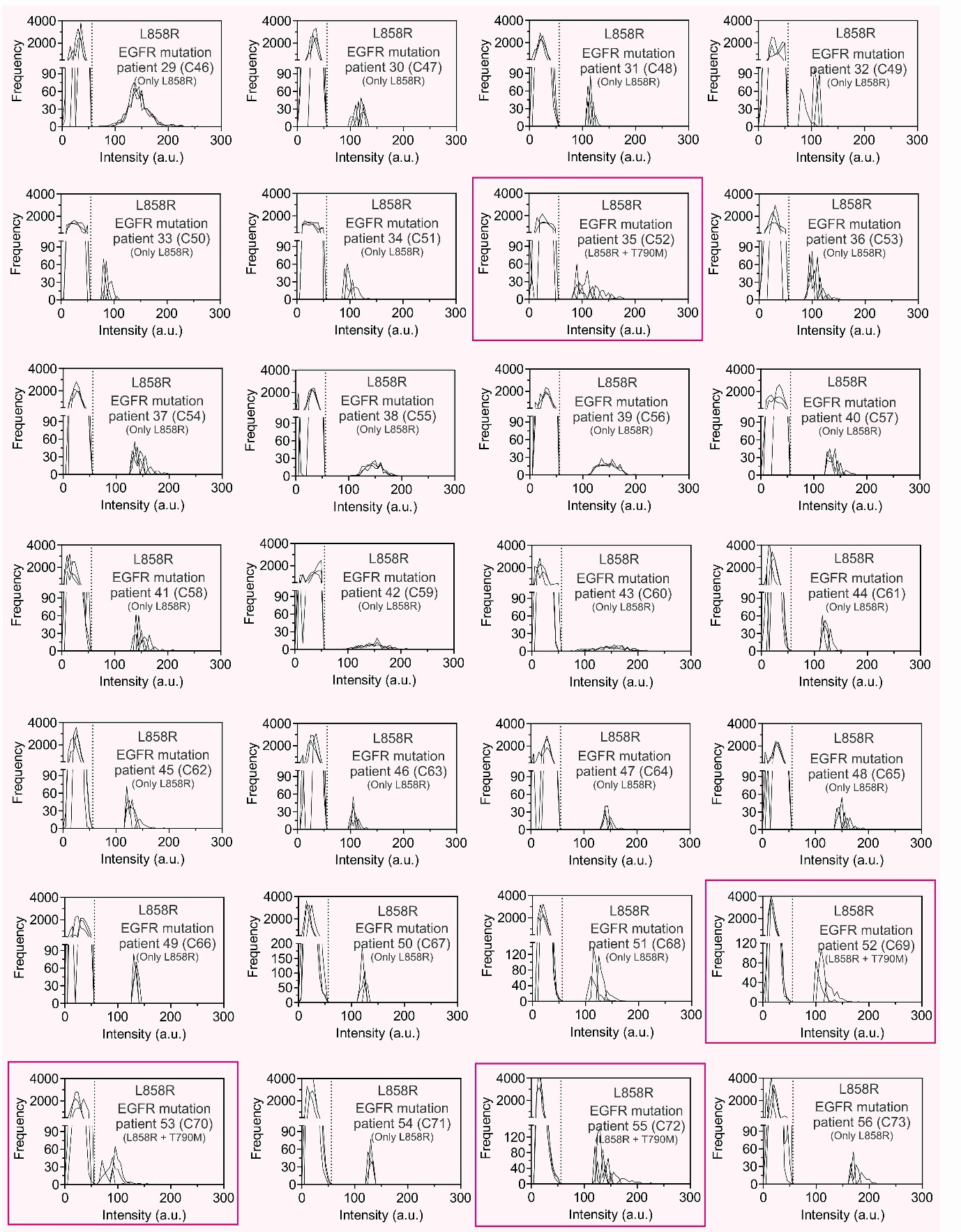

**Figure S21. Histogram of EV-CLIP droplets for L858R detection from clinical samples with EGFR mutation.** Histogram for intensity profile of L858R EV-CLIP droplets from 56 patients with lung cancer exhibiting EGFR mutations. There are peaks in which intensities are higher than the background signal, indicating the presence of L858R mRNA containing EVs within the droplet.

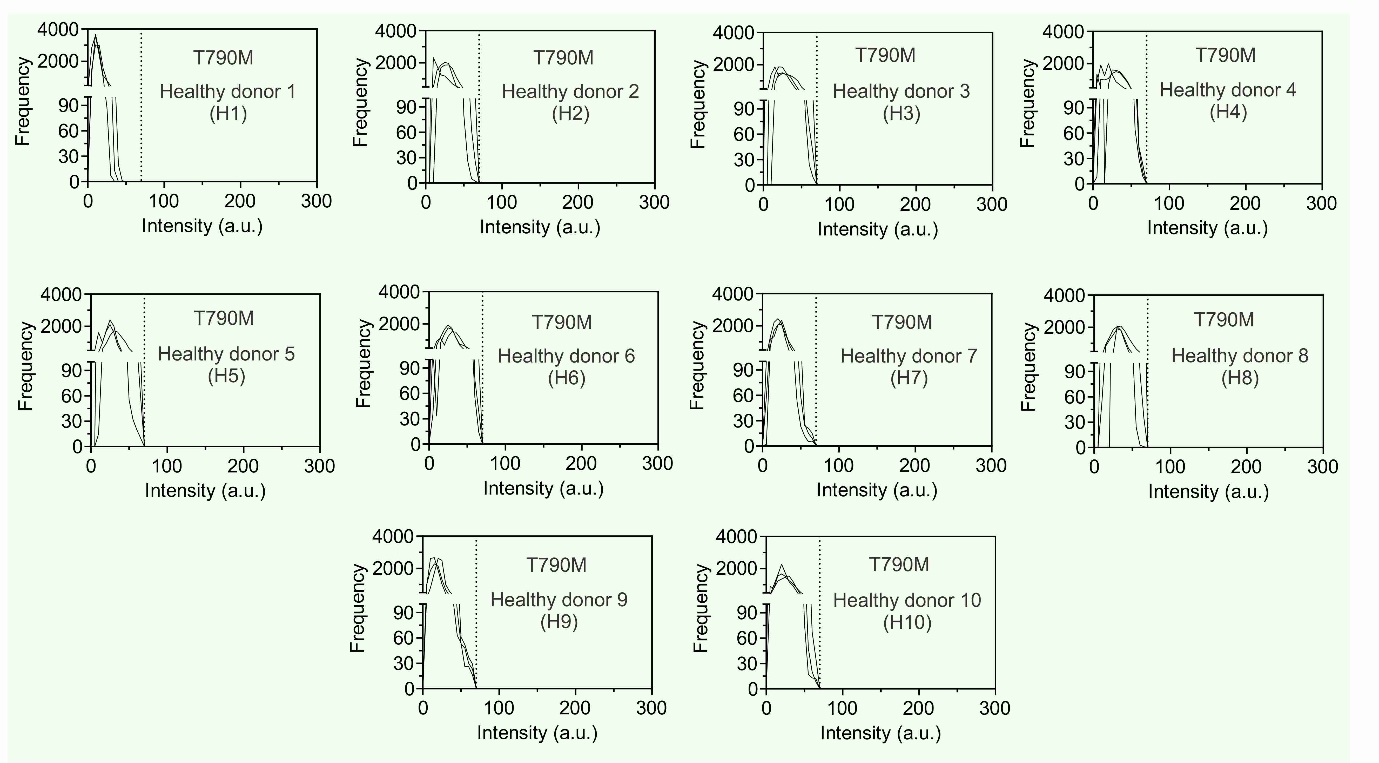

**Figure S22. Histogram of EV-CLIP droplets for T790M detection from healthy donor.** Histogram for intensity profile of T790M EV-CLIP droplets from 10 healthy donors.

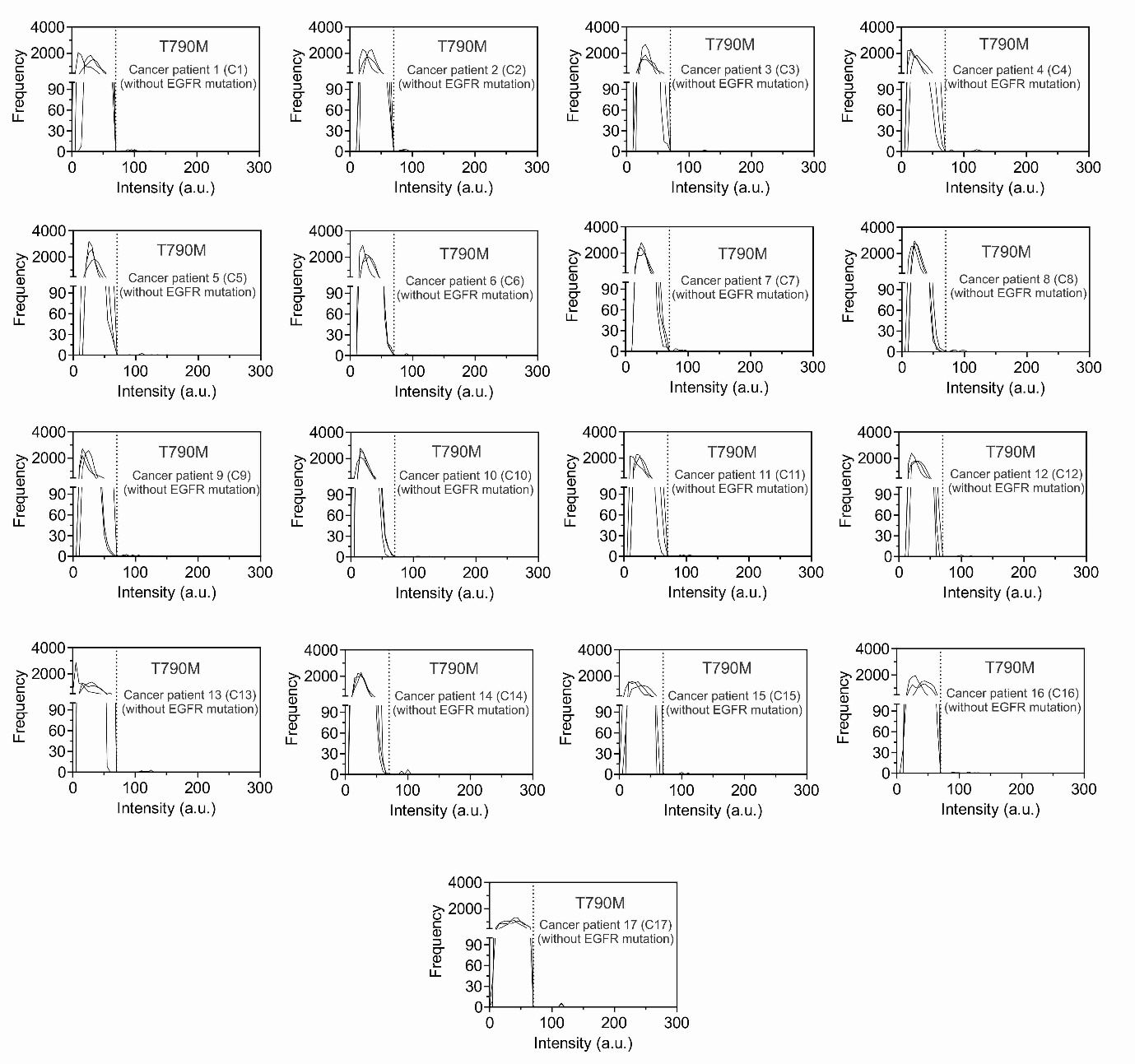

**Figure S23. Histogram of EV-CLIP droplets for T790M detection from clinical samples without EGFR mutation.** Histogram for intensity profile of T790M EV-CLIP droplets from 17 patients with lung cancer without EGFR mutations.

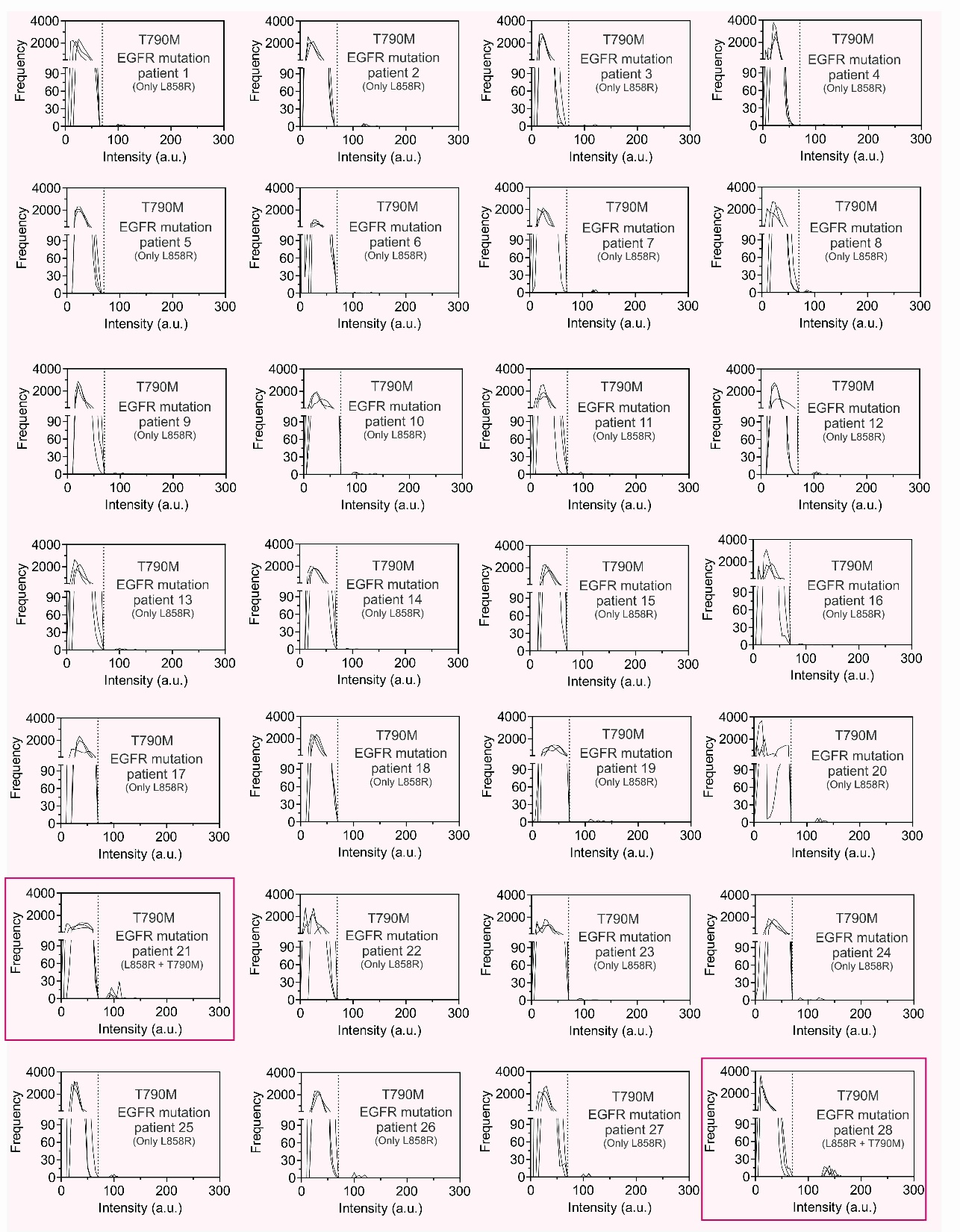

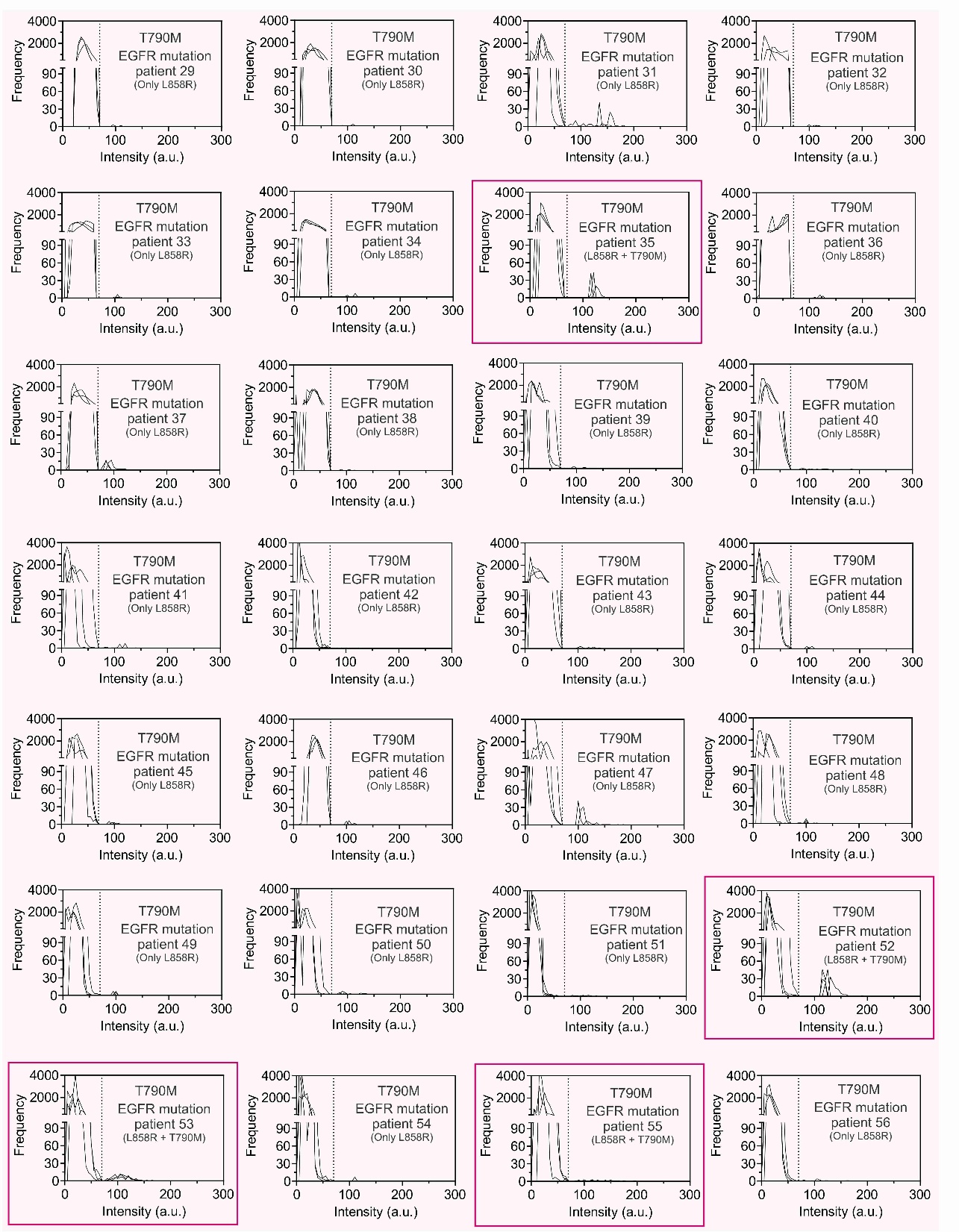

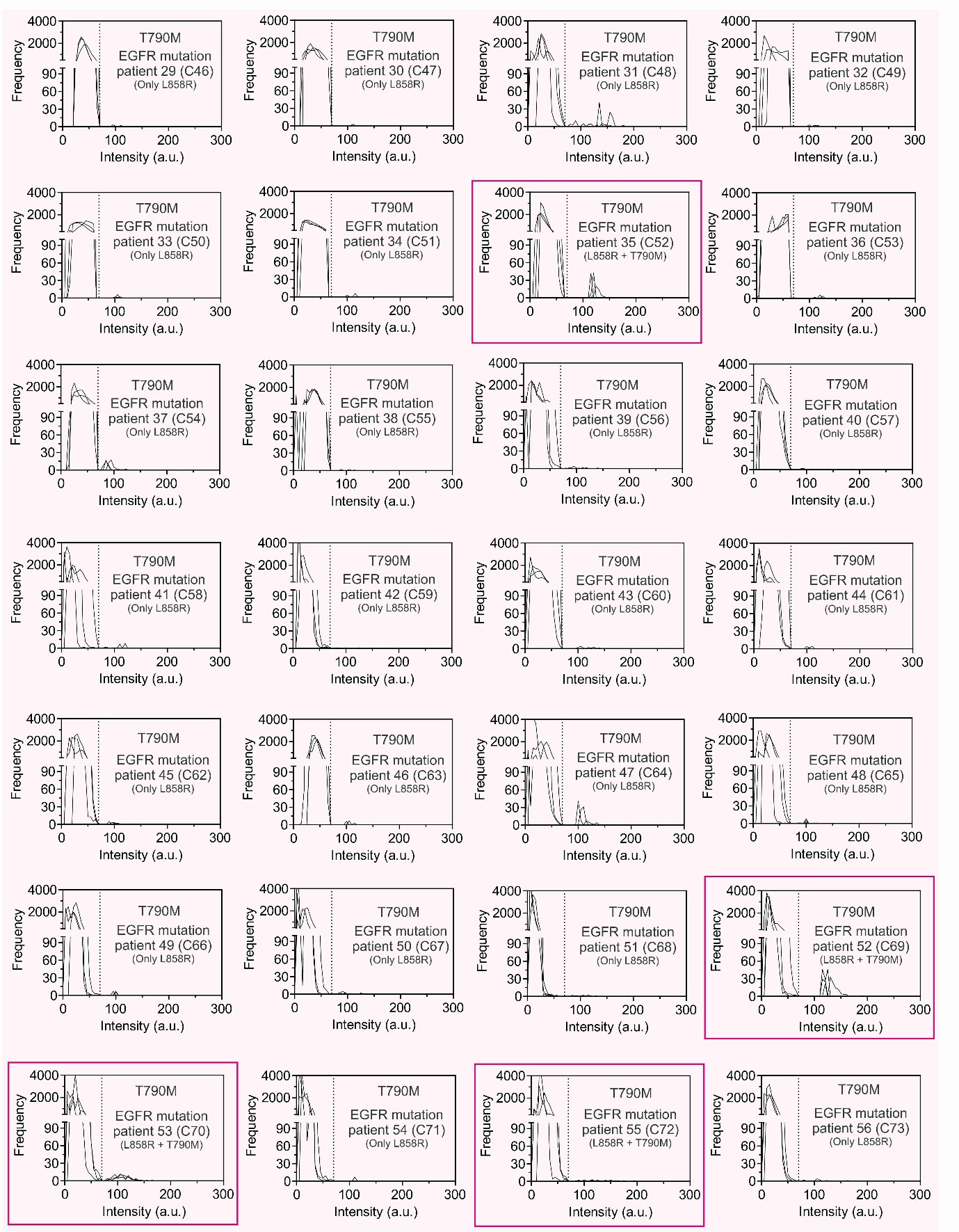

**Figure S24. Histogram of EV-CLIP droplets for T790M detection from clinical samples with EGFR mutation.** Histogram for intensity profile of T790M EV-CLIP droplets from 56 patients with lung cancer exhibiting EGFR mutations. There are peaks in which intensities are higher than the background signal only in the patients exhibiting T790M and in patient a, b, and c, as further elaborated in the main text, indicating the presence of T790M mRNA containing EVs within the droplets.

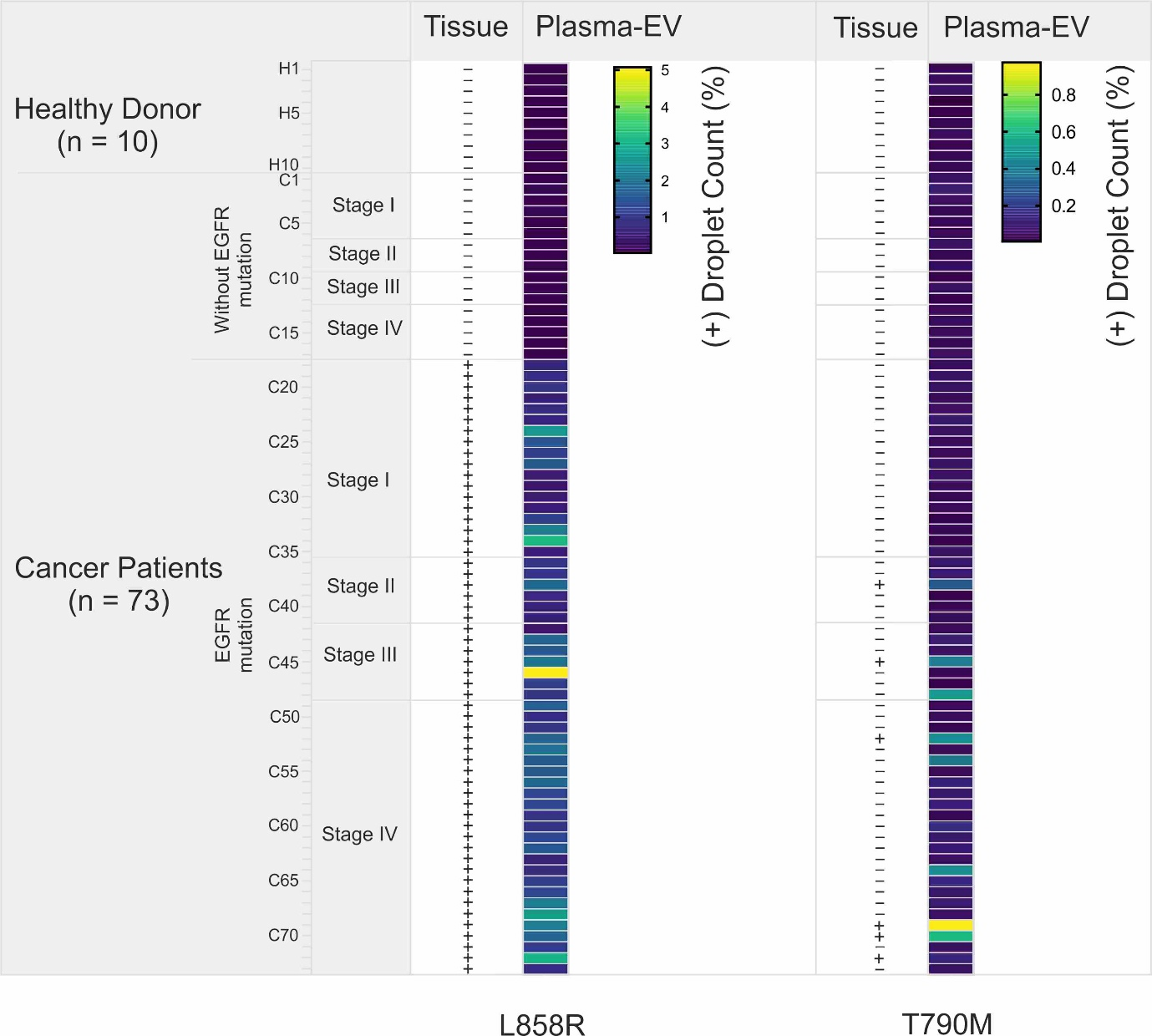

**Figure S25. Detection of EGFR L858R and T790M mutations from lung cancer samples using the EV-CLIP method.** A heatmap of 10 healthy donors and 73 patients with lung cancer for mutation detection.

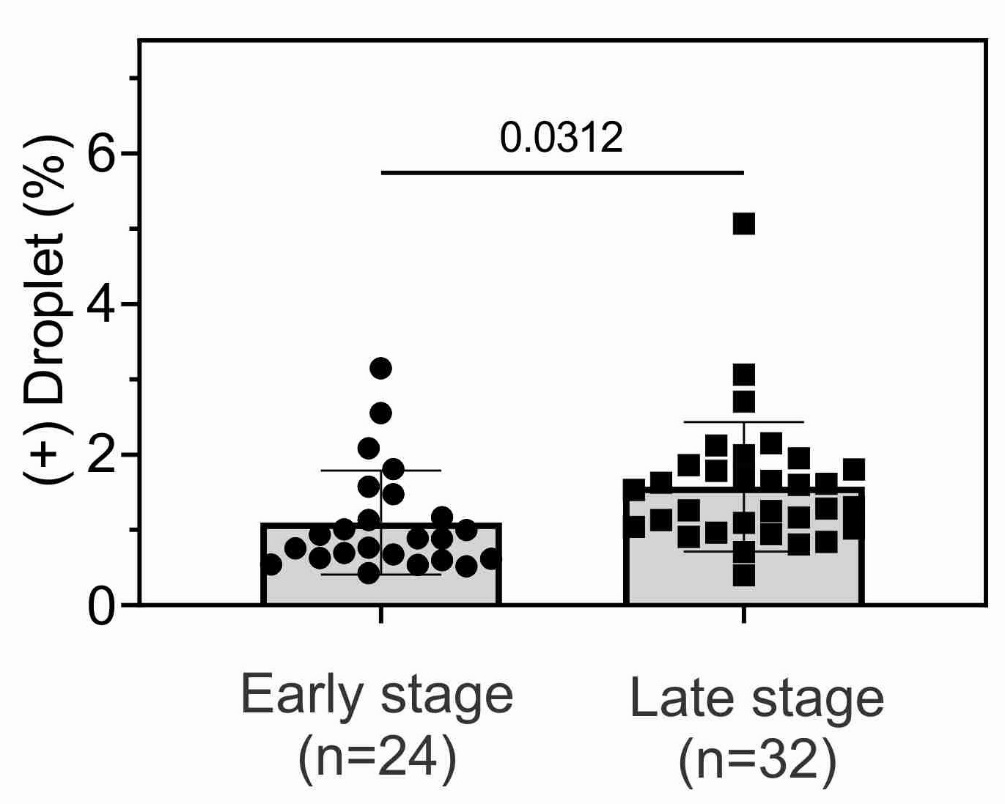

**Figure S26. Comparison of positive droplets % between patients with early and late-stage lung cancer.** Early-stage cancer comprises stages I and II, meanwhile late-stage cancer comprises stages III and IV. Each data point represents mean ± SD, n = 3. All statistical analyses were conducted using a two-tailed unpaired Student’s *t*-test.

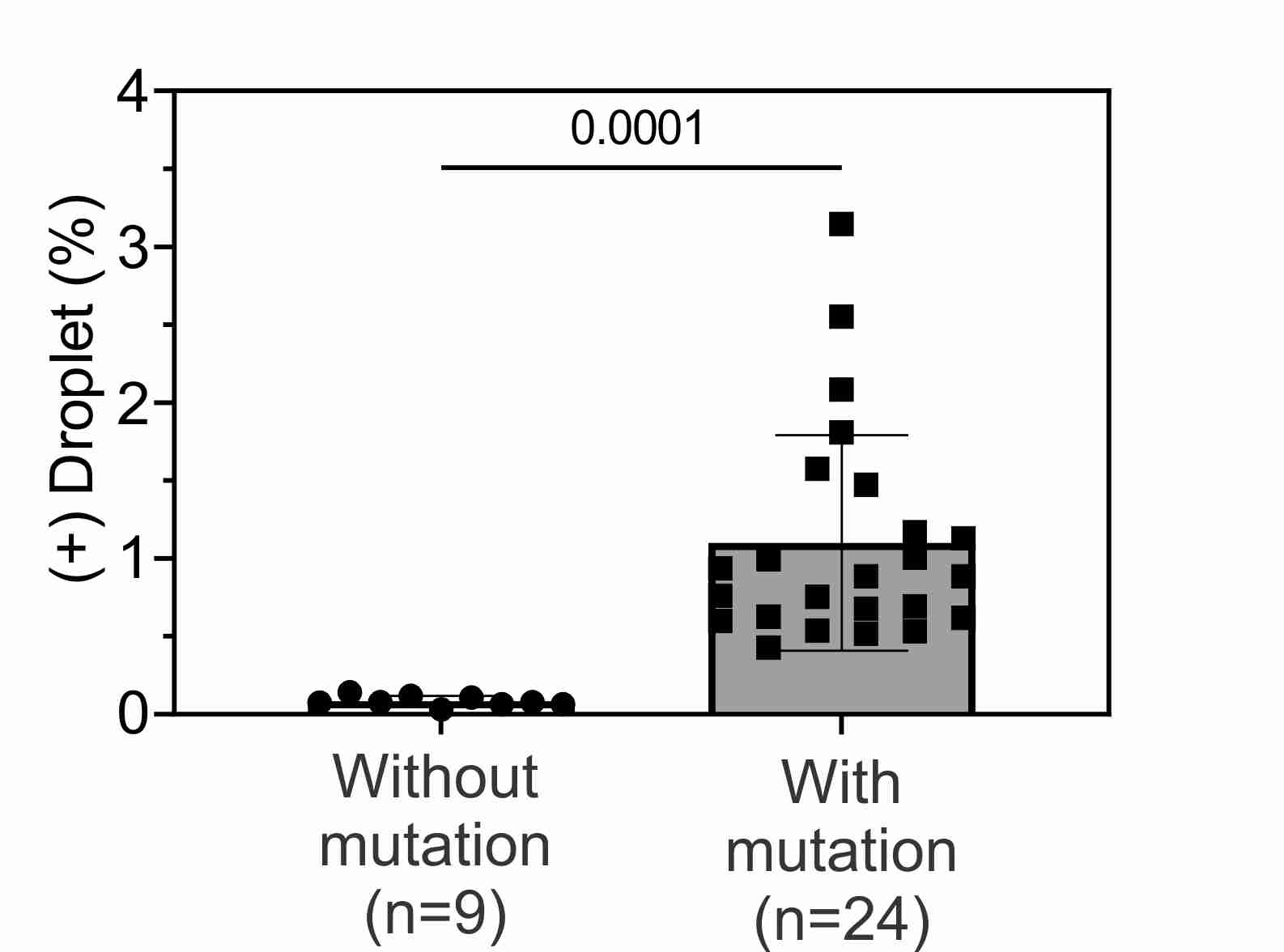

**Figure S27. Comparison of positive droplets % between patients with early-stage lung cancer with and without L858R mutation.** Early-stage cancer comprise stages I and II. Each data point represents mean ± SD, n = 3. Statistical analysis was conducted using a two-tailed unpaired Student’s *t*-test.

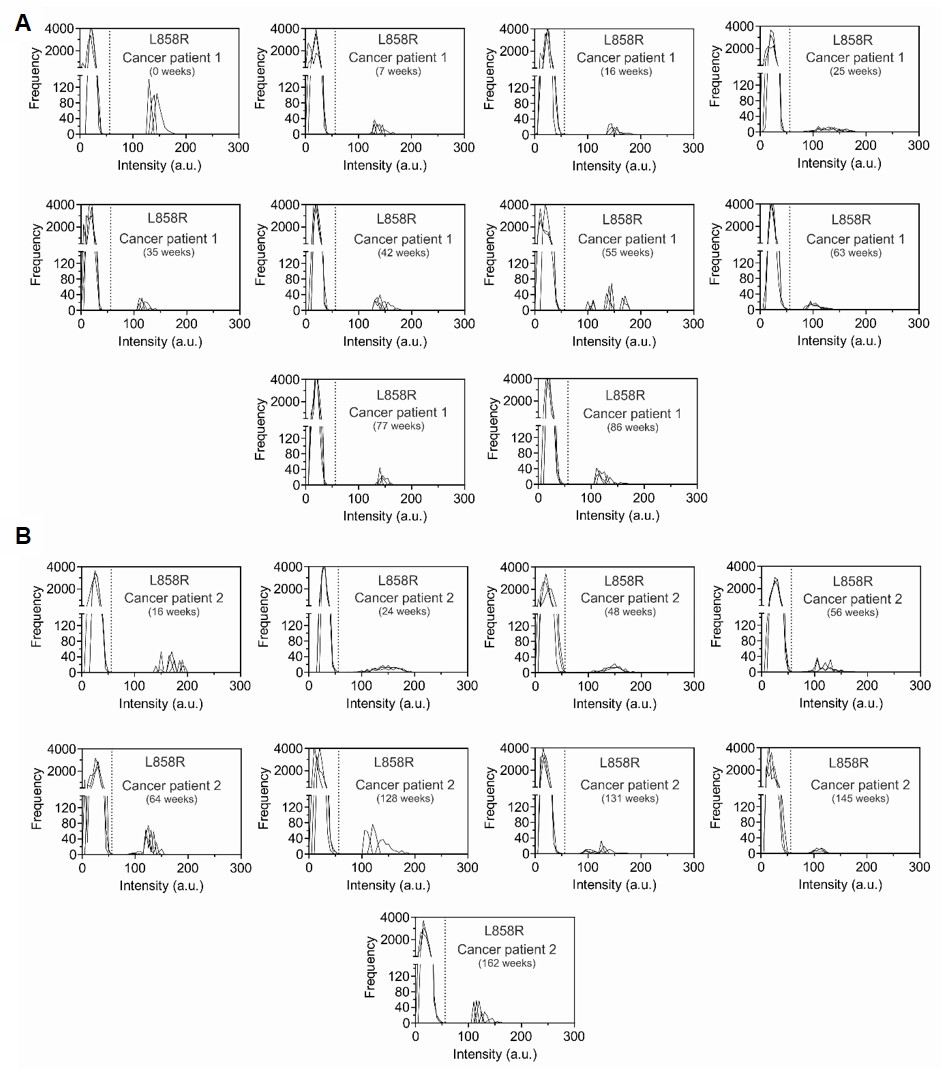

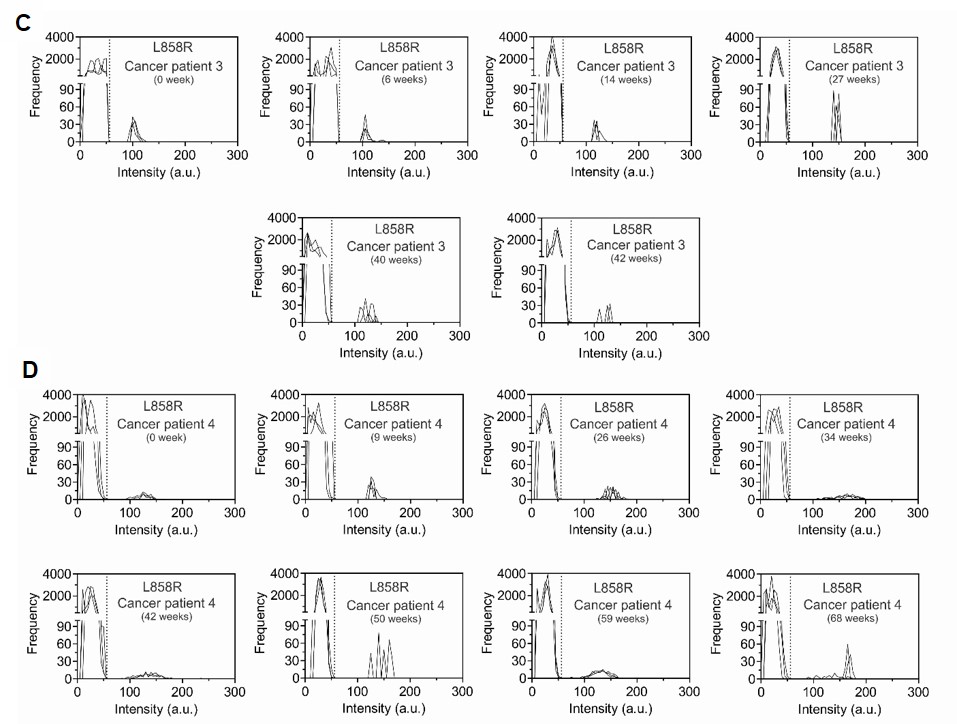

**Figure S28. Histogram of EV-CLIP droplets for longitudinal study of L858R mutation in patients with lung cancer.** Histogram of intensity profile for L858R EV-CLIP droplets from 33 samples of 4 patients with lung cancer exhibiting EGFR mutations, taken at various time points. (**A**) Histogram of intensity profile for plasma samples of patient 1, spanning 86 weeks. (**B**) Histogram of intensity profile for plasma samples from patient 2, spanning 162 weeks. (**C**) Histogram of intensity profile for plasma samples of patient 3, spanning 42 weeks. (**D**) Histogram of intensity profile for plasma samples of patient 4, spanning 68 weeks. There are peaks in which intensities are higher than the background signal, indicating the presence of L858R mRNA containing EVs within the droplets.

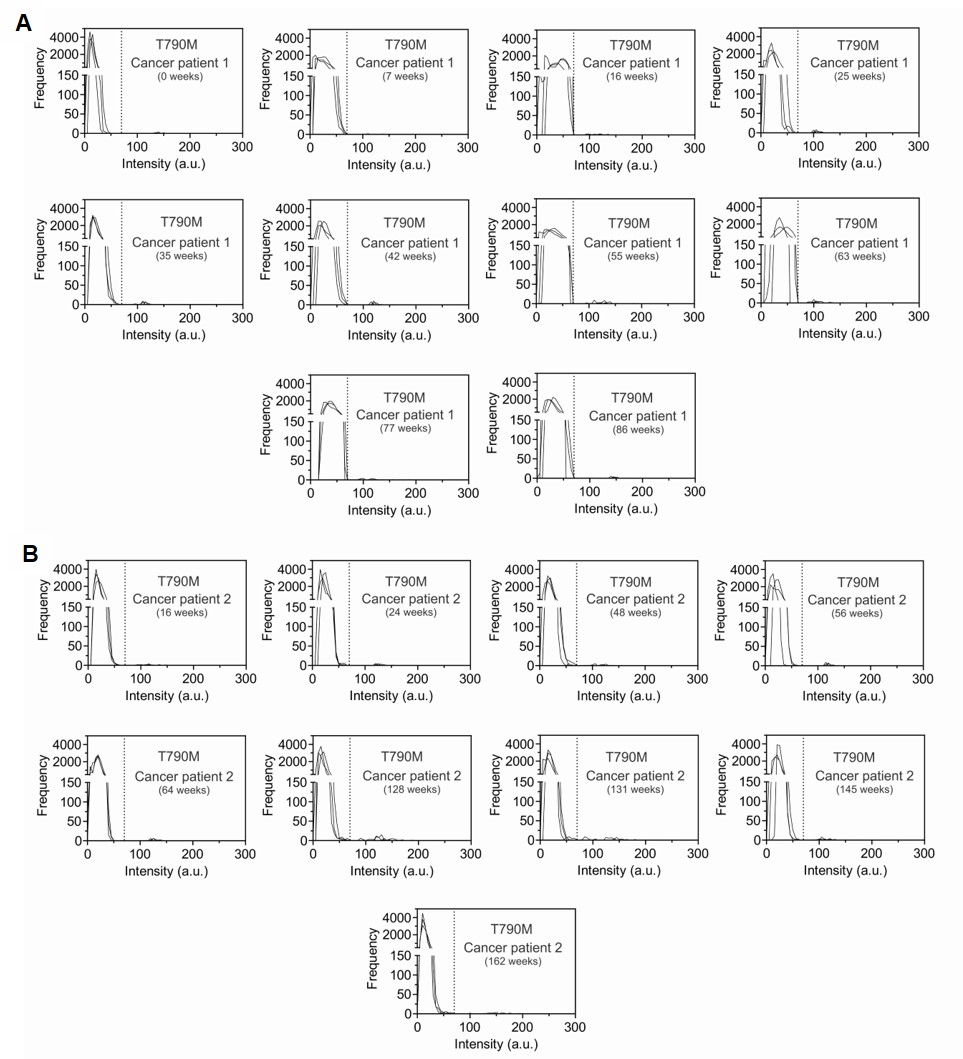

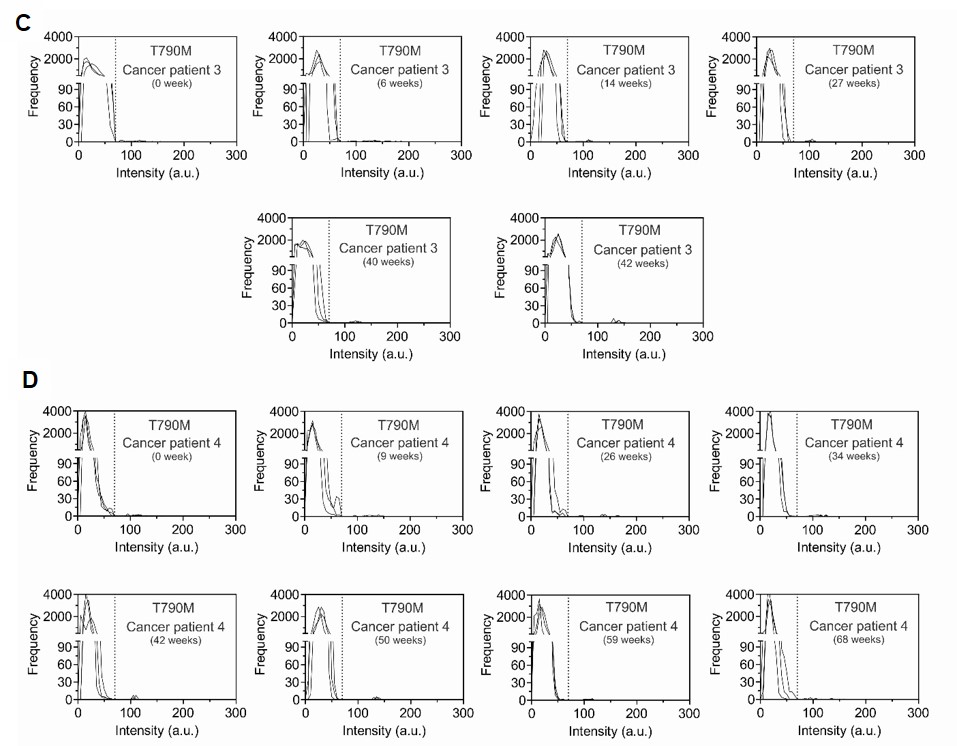

**Figure S29. Histogram of EV-CLIP droplets for longitudinal study of T790M mutation in patients with lung cancer.** Histogram of intensity profile for T790M EV-CLIP droplets from 33 samples of 4 patients with lung cancer exhibiting EGFR mutations, taken at various time points. (**A**) Histogram of intensity profile for plasma samples of patient 3, spanning 42 weeks. (**B**) Histogram of intensity profile for plasma samples from patient 4, spanning 68 weeks. (**C**) Histogram of intensity profile for plasma samples of patient 1, spanning 86 weeks. (**D**) Histogram of intensity profile for plasma samples from patient 2, spanning 162 weeks. There are peaks in which intensities are higher than the background signal, indicating the presence of T790M mRNA containing EVs within the droplets.

| miR-21 | TCAACA/iCy3/TCAGTCTGATAAGCTAGTATTATCAGACTGA/BHQ2 |
| --- | --- |
| EFGR (p.L858R mutation) | TTGGCC/iCy3/CGCCCAAAATCTGTGATTAGATTTGGGCG/BHQ2 |
| EFGR (p.T790M mutation) | AGCTGC/iCy5/ATGATGAGCTGCACGGTGGCAGCTCATCAT/BHQ2 |

**Table S1. Sequences used for the miR-21, EGFR L858R, and T790M mutations.**

The parts highlighted in red represent the fluorophore and quencher, respectively. These sequences were obtained from previously published study (*42*).

**Table S2.** Comparison between the EV-CLIP method and previously reported approaches for analyzing EV-derived miRNA. EV-CLIP method enhances EV-derived RNA detection sensitivity, requires < 20 µL plasma sample, and eliminates the need for sample preparation, enabling rapid digital detection.

| Detection Method | Bulk or Digital | Sample | EV isolation | miRNA  Biomarker | Time^1^ | LOD | Ref |
| --- | --- | --- | --- | --- | --- | --- | --- |
| CLSM^2^ imaging of EV fusion with virus-mimicking vesicles containing MB^3^ | Bulk | Serum | No | miR-21 | 2 h | 1.3 nM of miR-21 | (*31*) |
| TIRF imaging after delivery of split DNAzyme probe on EVs treated with streptolysin O | Bulk^2^ | Serum | No | miR-21 | 1 h | 378 copies of miR-21/µL | (*52*) |
| TIRF^4^ imaging of EV fusion with tethered cationic lipoplex nanoparticles (tCLN) | Bulk | EVs isolated from 60 µL serum | Yes | miR-21, miR-25, miR-155, miR-210, miR-486 | EV prep + 2 h | NA | (*53*) |
| EV fusion with CLP-MBs^5^ in a micro-mixer chip, followed by fluorescence detection using a microplate reader | Bulk | EVs isolated from 30 µL serum | Yes | miR-21 | EV prep. + 10 min | 2.06 × 10^9^ EVs/mL  (3.4 pM) | (*54*) |
| EV fusion with MB-containing vesicles modified with zipper DNA constructs followed by fluorescence intensity detection using a microplate reader | Bulk | EVs isolated from 5 mL plasma | Yes | miR-21 | EV prep. + 30 min | 5 x 10^8^ EVs/mL  (0.8 pM) | (*55*) |
| Flow cytometry analysis after aptamer-mediated fusion with MB-encapsulated liposomes | Bulk | 5 µL plasma | No | miR-21 | 30 min | 0.14 μg/mL EVs  (7.4 fM EV miR-21) | (*32*) |
| Surface plasmon-enhanced fluorescence spectroscopy detection of EV surface proteins and microRNA pairs | Bulk | EV from 50 µL serum | Yes | miR-21 | 4 h | 5 x 10^9^ EVs/mL  (8.31 pM) | (*56*) |
| EV fusion with liposomes containing Cas13 complex | Bulk | 500 µL of diluted plasma | No | miR-21 | 1 h | 1.2 x 10^3^ EVs/mL (2.0 aM)  2.14x10^3^ EVs/mL in plasma (3.6 aM) | (*57*) |
| Digital detection after charge-based fusion of EVs with liposomes containing MB | Digital | 20 µL Plasma | No | miR-21 | 20 min | 79 EVs/µL (0.13 fM) | This Work |

^1^ Assay time includes the incubation before measurement but does not account for device preparation or sample preparation steps; ^2^confocal laser scanning microscopy (CLSM); ^3^molecular beacon (MB); ^4^total internal reflection fluorescence (TIRF); ^5^cationic lipoplexes containing molecular beacons (CLP-MBs)

**Table S3.** Comparison between the EV-CLIP method and previously reported approaches for analyzing EV-derived mRNA.

| Detection Method | Bulk or Digital | Sample | EV isolation | RNA isolation | Amplification | mRNA  marker | Time^1^ | LOD | Ref |
| --- | --- | --- | --- | --- | --- | --- | --- | --- | --- |
| TIRF imaging of EV fused with tethered LPHNs^2^ containing CHDC^3^ | Bulk | Serum  (10 µL) | No | No | Signal Amplification | *GPC1* | 2 h | 60 EVs per µL^4^  (~ 0.1 fM) | (*58*) |
| TIRF imaging of EV fusion encapsulated in CLP^5^ tethered on a gold-coated glass slide chip | Bulk | Serum  (20 µL) | No | No | - | *KRAS*  *EGFR* | 2 h | NA | (*42*) |
| Three-way junction-based isothermal gene amplification | Bulk | mRNA (10 ng) | Yes | Yes | Isothermal gene amplification | *PPP1R1B* | RNA prep + 30 min | NA^6^ | (*59*) |
| Chip-based detection using 3D nanostructured hydrogels with catalytic hairpin assembly | Bulk | EVs isolated from 400 µL plasma | Yes | No |  | *ERBB2, GAPDH* | EV prep + 2 h | NA^7^ | (*60*) |
| Immuno-magnetic exosome RNA analysis coupled with real-time PCR | Bulk | mRNA isolated from 100 µL plasma | Yes | Yes | Real-time PCR | *EPHA2, EGFR, PDPN, MGMT, APNG* | 2 h + RNA prep | NA | (*61*) |
| Droplet digital PCR | Digital | mRNA isolated from 2 mL plasma | Yes | Yes | ddPCR | EGFR*vIII* | RNA prep + ~40 min | 5.53 copies of *EGFRvIII* | (*62*) |
| Digital detection after charge-based fusion of EVs with liposomes containing MB | Digital | Plasma  (20 µL) | No | No | No | *EGFR* | 20 min | L858R:  1348 EVs/µL (2.24 fM)  T790M:  2696 EVs/µL (4.48 fM) | This work |

^1^ Assay time includes the incubation before measurement but does not account for device preparation, sample preparation steps such as EV isolation or RNA extraction, and amplification; ^2^lipid-polymer hybrid nanoparticles (LPHNs); ^3^catalyzed hairpin DNA circuit (CHDC); ^4^ LOD calculated from the intensity of GPC1 mRNA from AsPC-1 cell line-derived EVs; ^5^cationic lipoplex nanoparticles (CLP); ^5^overhang molecular beacon with internal dye (Ohi-MBs); ^6^LOD of 1.23 pM was estimated from PP1R1B RNA analysis; ^7^LOD of 58.3 fM was estimated from ERBB2 RNA detection.

**Table S4.** Comparison between the EV-CLIP method and previously reported digital detection approaches for analyzing EV proteins.

| Detection Method | Biomarker | | LOD | Ref |
| --- | --- | --- | --- | --- |
| Surface-anchored nucleic acid amplification and compartmentalization in droplet | Protein | GPC-1 | Not mentioned | (*63*) |
| Single exosome counting enzyme-linked immunoassay | Protein | GPC-1 | 10 enzymes labeled EV complexes/µL (0.02 fM) | (*64*) |
| Droplet microfluidics with magnetic beads with antibody-coated with DNA with PCR | Protein | EGFR, EPCAM, GZMB, TCF7 and PD-L1 | 38 EVs/µL (0.06 fM) | (*65*) |
| Antibody-based immune sequencing method in droplet | Protein | CD9, CD81, CD63, CD11b, F4/80 and CD45 | Not mentioned | (*66*) |
| High throughput droplet digital enzyme-linked immunosorbent assay | Protein | CD81 | 9 EVs/µL (0.01 fM) | (*67*) |
| Digital detection after charge-based fusion of EVs with liposomes containing MB | Nucleic Acid  (miRNA, mRNA) | miR-21  EGFR L858R, T790M | miR-21: 79 EVs/µL (0.13 fM)  L858R: 1348 EVs/µL (2.24 fM)  T790M: 2696 EVs/µL (4.48 fM) | This Work |

**Table S5.** **Patient information.** Basic information, including gender, age, histology types, smoking status, mutation, and stage of 83 patients enrolled in the experiment.

| Characteristics | Healthy (n=10) | Patients without EGFR mutation (n=17) | Patients with EGFR mutation (n=56) |
| --- | --- | --- | --- |
| Gender |  |  |  |
| Female | 5 | 9 | 32 |
| Male | 5 | 8 | 24 |
| Age |  |  |  |
| >60 | 4 | 16 | 8 |
| ≤60 | 6 | 1 | 48 |
| Histology types |  |  |  |
| Adenocarcinoma | 0 | 17 | 53 |
| Non-adenocarcinoma | 0 | 0 | 3 |
| Smoking status |  |  |  |
| Nonsmoker | 7 | 0 | 18 |
| Smoker | 3 | 0 | 11 |
| Unknown | 0 | 17 | 27 |
| Mutation |  |  |  |
| L858R |  | 0 | 50 |
| L858R+T790M |  | 0 | 6 |
| Stage |  |  |  |
| I | 0 | 6 | 18 |
| II | 0 | 3 | 6 |
| III | 0 | 3 | 7 |
| IV | 0 | 5 | 25 |
